# Development and characterization of a scalable calcium imaging assay using human iPSC-derived neurons

**DOI:** 10.1101/2025.09.05.674208

**Authors:** Martin Dietrich Haustein, Caroline Deymier, Simon Schlienger, Laurane Lexcellent-Bissler, Jacques Mawet, Eric Gutknecht, Laure Thenoz, Pijus Brazauskas, Bérengère Renault, Stéphanie Brun, Simone Winistörfer, Thomas Portmann

## Abstract

Neuroscience drug discovery is challenged by the brain’s structural and cell-type complexity, which is difficult to model in cellular systems compatible with high-throughput screening methods. Calcium oscillation assays, that harness neurons’ intrinsic capability to develop functional neural networks in cell culture, are currently the closest cellular models with a relevant functional endpoint to model human neuronal circuitry in a dish. Here we further developed this useful assay towards scalable drug discovery applications. We show the importance of defined neuron-to-astrocyte ratios for optimal cellular distribution and surface adherence in HTS-compatible cell culture vessels and how the cell type ratios affect network firing patterns. Increasing the neuron density resulted in decreased network spike frequencies, but increased network spike amplitudes. We identified DAPT, a molecule previously shown to promote neuronal maturation and synapse formation, as a negative regulator of astrocyte viability. Furthermore, inclusion of GABAergic neurons in the cocultures increased the network spike frequency while reducing network spike amplitudes. The GABA A receptor antagonist bicuculline did not affect network spike frequency, but increased network spike amplitudes. In order to access local field activity in an automated and scalable calcium imaging environment, we developed a pixel-based analysis for plate reader data. This method revealed that the effect of GABAergic neurons and bicuculline was restricted to local field calcium activity that coincided with synchronized network spikes. Our observations are consistent with previous findings suggesting that the presence of GABAergic neurons decreases synchronization and network spike participation of local neuronal activity, thus potentially echoing aspects of GABA action in vivo, and dysregulation thereof in pathological conditions.

## 1 Introduction

Drug discovery efforts to address pathological conditions of the central nervous system (CNS) have historically suffered from a lack of clinically translatable in vitro systems that are useful both for high-throughput screening (HTS) and efficient downstream evaluation of candidate molecules. While the integration of single-cell resolution technologies in neuroscience research has advanced our understanding of pathological mechanisms in the neural circuitry and in specific neuronal and glial cell types, modeling these mechanisms in cell culture systems has remained a challenge. This is due mainly to the complexity of the CNS architecture, with a myriad of different cell types interacting in health and disease, as well as due to the difficulty of generating precise replica cell types of translational relevance in a cell culture dish. The latter is of particular importance as some of the most promising drug targets in the CNS, including G-protein coupled receptors, are found to be expressed in unique and highly cell type-selective patterns. Hence, replicating (1) the precise CNS cell type(s) of interest for drug treatment in patients and (2) their endogenous target expression including the surrounding protein signaling interactome is of utmost importance to properly assess the translational potential of candidate compounds early in the CNS drug discovery process.

The discovery of somatic reprogramming and the development of protocols for differentiation of human induced pluripotent stem cells (iPSC) into neurons has helped the field to overcome major limitations imposed by the inaccessibility of live human CNS tissue for research (Takahashi et al., 2007). Importantly, human iPSC offered new opportunities to complement rodent CNS-derived in vitro models and heterologous expression systems with human CNS cell types in drug discovery research. While early iPSC-derived CNS models were generated by neuronal differentiation protocols taking embryonic brain development as template (Brennand et al., 2011; Paşca et al., 2011; Shcheglovitov et al., 2013), cellular reprogramming by transcription factor over-expression is increasingly adopted for properties that are easier reconciled with scalability in drug discovery assays (Saavedra et al., 2021; Yang et al., 2017; Zhang et al., 2013). These properties include the rapid and relatively homogeneous generation of specific neuron or glia types within days. Scaled production and cryopreservation of reprogrammed CNS cell types further allow for rapid integration into assays without the need of starting from pluripotent stem cells. Essentially, the focus is on generating cell types “close enough” to their in vivo counterparts with regards to target expression and key functional properties. And this, achievable with relatively simple, rapid, and robust cell culture protocols. Transcription factor-based reprogramming allows for comparably efficient generation of highly pure human in vitro CNS-like cell types. Combinations of specific transcriptions factors, further termed herein reprogramming codes, allow for generation of induced neurons, including glutamatergic excitatory neurons (GLUT iN, reprogramming code: Ngn2)(Zhang et al., 2013), GABAergic inhibitory neurons (GABA iN: Ascl1 and Dlx2)(Yang et al., 2017), and various glia types that resemble key CNS cell types of the human brain.

A mainstay of CNS drug discovery and development is the calcium oscillation assay, which allows for rapid assessment of key functional properties of neural networks and their modulation by therapeutic candidate molecules. In this assay, synchronized firing of an entire neural network results in globally oscillating calcium signals that can be measured on plate readers based on fluorescent signals from calcium-sensitive dyes or genetically encoded calcium indicators. For development of synchronized neural network activity of human GLUT iN, astrocyte-derived factors to support synapse formation and maturation are key. The simplest way of providing these factors is by culturing neurons in astrocyte-conditioned medium (Shan et al., 2024). The alternative is co-culturing neurons with astrocytes in order to allow for astrocytes to provide real-time metabolic support and actively modulate synapse development. Rodent astrocytes from fetal or neonate brain have widely been used for ease of access (Patzke et al., 2015). Primary fetal astrocytes (pA) of human origin in turn allow for development of oscillation assays of completely human cellular origin (Bassil et al., 2021; Saavedra et al., 2021). However, while many versions of calcium oscillation assays using human neurons have been presented, few only did so within a scalable environment. For example, multi-week protocols for human neuron-based calcium oscillation assays often still include use of low-throughput surfaces, such as glass coverslips (Sun & Südhof, 2021). With few exceptions, the precise ratio of neurons and astrocytes for optimal network development, and how it affects oscillation characteristics has not been described in detail, and finally, key cell culture properties including cell attachment and cellular distribution in 2D (versus aggregate formation) and their impact on calcium signals remain underexplored. Also, the importance of GABAergic inhibition, important for control and synchronization of neural network activity in vivo and playing an important role in therapeutic action of psychiatric and anti-epileptic drugs, merits further exploration in the context of all-human in vitro neural network models.

Here, we set out to test various parameters of a human GLUT iN-based calcium oscillation assay, towards scalable applications including drug screening on fully functional neural networks. We characterized key parameters to allow for implementation in a standard high-throughput environment, including culture on HTS-capable optical cell culture plates, and the impact of neuron-astrocyte ratios on firing properties. We find that neuron-astrocyte ratios influence not only network firing properties, but also cellular attachment and the tendency of GLUT iN to form clusters or remain evenly distributed over time. We show that brief treatment with DAPT, a gamma secretase inhibitor, changes the ratio of neurons and astrocytes by selectively decreasing the viability of astrocytes. And finally, we describe how addition of GABA iN and pharmacological inhibition of GABA A receptors affect network firing properties and synchronization of local field activity across the well surface, as demonstrated by a novel, pixel-resolution analysis of plate reader data.

## 2 Materials and methods

### 2.1 Human iPSC lines and primary astrocytes

The human iPSC lines and primary fetal astrocytes used in this study were obtained from commercial sources (StemRNA Human iPSC 771-3G from Reprocell, and Human Astrocytes from ScienCell) and public repositories (EBiSC: BIONi010-C-13 iPSC line).

### 2.2 Human iPSC culture and engineering

Details of human iPSC culture and engineering are described in the supplementary methods. Manufacturers, reagent item numbers, and PCR primer sequences are provided in Table S1. Briefly, human iPSC were cultured in mTeSR Plus on Geltrex-coated cell culture vessels using standard cell culture conditions and passaged using ReLeSR according to manufacturers’ recommendations.

For CRISPR-based genome engineering, human iPSC were electroporated with ribonucleoprotein complexes and 1.5 µg donor DNA constructs using a Neon electroporator and a single 30 ms pulse at 1100 V. For recovery and to increase the frequency of homology-directed repair (HDR), cells were kept at 32 °C in the presence of CloneR2 and HDR-Enhancer V1, an inhibitor of non-homologous end joining (NHEJ). After 2 days, the medium was replaced with mTeSR Plus supplemented with the appropriate selection antibiotic (Puromycin: 0.3 µg/mL / Blasticidin: 2 µg/mL). For Cre recombinase-mediated cassette exchange (RMCE), cells were pulsed with doxycycline for 24 h prior to electroporation. Cells were electroporated with 1.5 µg RMCE donor plasmid using a triple 20 ms pulse at 1200 V. Cells were recovered in the presence of CloneR2. After 2 days, the medium was replaced with mTeSR Plus supplemented with 2 µg/mL Blasticidin. Clonal selection was either performed by limiting dilution in 96-well plates or by sparce single cell seeding in 10 cm dishes and manual colony picking. Standard PCR genotyping was used to verify all genomic engineering steps. All iPSC lines were checked by karyotyping, and the master iPSC and Bi-GCaMP6f-2A-RFP lines were further checked for microdeletions or microduplications by array comparative genomic hybridization. Pluripotency marker staining was occasionally included to spot-check pluripotency state of the cell (Figure S4D).

Schematic views of DNA donor constructs are shown for homology-directed repair (HDR) after CRISPR (Figure S1A, Figure S4A, C) and for RMCE (Figure 3A). For HDR constructs, the selection cassettes were flanked by compatible Lox sites (LoxFas and LoxN) to enable removal by Cre recombinase. A Cre recombinase coding sequence was coupled to EGFP via a P2A sequence and put under control of a tetracycline responsive Tet-ON promoter. A reverse Tet transactivator (rtTA) coding sequence was put under control of the constitutive CAGGS promoter (Hitoshi et al., 1991). Selection cassettes flanked by compatible Lox sites were removed in the final master iPSC line by brief exposure to doxycycline. The Cre transgene was flanked by incompatible Lox sites (LoxP and Lox2272) to enable integration of any desired reprogramming code by RMCE. RMCE donor constructs contained the reprogramming code composed of the transcription factors ASCL1 and DLX2 (coupled by a 2A sequence) under control of a Tet-ON promoter. The herein presented design of iPSC lines served the sole purpose of assay feasibility evaluation, and any future use in, for example, drug discovery research may require further optimization.

### 2.3 Neural reprogramming of human iPSC and neuron-astrocyte coculture

Human iPSCs were seeded as single-cell suspension in the presence of 2 µM Thiazovivin on Matrigel coated culture vessels at a density of 10000 cells/cm^2^. From d1 to d3 the medium was exchanged to reprogramming medium (DMEM/F12 supplemented with N2, NEAA, 10 ng/mL hBDNF, 10 ng/mL hNT-3, 200 ng/mL mouse laminin, and 2 µg/mL doxycycline) and renewed daily. On d4 the young neurons were either cryopreserved in Bambanker or replated in neuronal medium (Neurobasal-A supplemented with B27, GlutaMAX, 10 ng/mL hBDNF, 10 ng/mL hNT-3, 200 ng/mL mouse laminin, and 2 µg/mL doxycycline) for coculture with astrocytes.

On d2 of reprogramming, human primary astrocytes were seeded on laminin-coated cell culture plates in Astrocyte medium. On d4, young neurons were seeded in neuronal medium on top. Half-media changes were performed every 2-3 days. From d7-d10, the neuronal medium was supplemented with 2 µM Ara-C and from d11 onwards with 2.5% FBS.

### 2.4 Cell viability assays

LDH assays were performed with the CytoTox 96 non-radioactive assay, according to manufacturer’s protocols. Briefly, cell culture conditioned medium was collected, spun down, mixed with CytoTox 96 Reagent, and incubated for 30 minutes at room temperature protected from light. The reaction was stopped by adding Stop Solution, and absorbance was measured at 490 nm with a SynergyMX microplate reader. Cell culture medium served as control. Data processing and analysis were performed according to manufacturer’s recommendations.

Realtime monitoring of pA apoptosis and proliferation was performed by seeding 10’000 pA/well in a laminin-coated 96 well-plate in Astrocyte medium. After 2 days, the medium was replaced by neuronal medium as used in the neuron-astrocyte coculture and supplemented with 5µM Incucyte® Caspase-3/7 Green Dye. Phase and fluorescent images were acquired with the Incucyte SX5 every 2h. Masks for the cell confluence and the dye fluorescence were directly generated in the Incucyte software.

### 2.5 Single-Nucleus RNA Sequencing (snRNA-seq) Library Preparation and Sequencing

For nuclei isolation from a 384-well plate, cells were lysed on ice using cold Lysis buffer (1x PBS, Tris-HCl 10 mM, NaCl 10 mM, MgCl_2_ 3 mM, NP40 0.025%, pH 7.4) for 3 min with 3-5 times gently mixing using a micropipette. Lysis was stopped by addition of cold Nuclei Wash and Resuspension Buffer (NWR Buffer: 1x PBS, 1% BSA, RNase Inhibitor 0.2 U/μL). The nuclei were collected by centrifugation, and washed twice with cold NWR buffer, before being resuspended in 1x PBS containing 0.04% BSA.

The 3’ CellPlex Kit Set A was used for labeling of nuclei with Cell Multiplexing Oligos (CMO) following closely the manufacturer’s recommendations (see supplementary methods for details). Nuclei concentration was measured using a Luna-FX7 automated cell counter, and diluted to 1500 cells/µL for subsequent snRNA-seq experiment.

Single-nucleus RNA-seq libraries were prepared using the Chromium Next GEM Single Cell 3ʹ Reagent Kit v3.1 (10x Genomics) according to the manufacturer’s protocol (CG000388). Briefly, single nuclei were captured, barcoded, and reverse-transcribed, followed by cDNA amplification and library construction. Library size distribution was assessed using Fragment Analyzer NGS High Sensitivity Kit (Agilent Technologies), and library concentrations were quantified using KAPA Library Quantification Kit (Roche Diagnostics) according to the manufacturer’s instructions. Cell Multiplexing Oligo (CMO) libraries and gene expression libraries were pooled at a 1:4 molar ratio and sequenced on an Illumina NextSeq 2000 platform using a P2 100-cycle kit (Illumina).

Sequencing reads were aligned to the human reference genome (GRCh38; GENCODE v32/Ensembl 98) using the Kallisto bustools (kb-python) pipeline with the -nac parameter to obtain single-cell gene count matrices (Melsted et al., 2021). CMO assignment was performed using Cell Ranger v7.2.0 (10x Genomics).

For cell type annotation and clustering, cell type specific genes were identified by correlation to known cell type-specific marker genes. In addition, cell type-specific genes were confirmed based on the Jensen-Shannon divergence of their expression distribution from the expression distribution of a hypothetical, perfectly specific gene (Lin, 1991). Graphing was done using Python as described under Statistical analysis and graphing.

### 2.6 Calcium imaging

Calcium imaging was conducted at d28-31 as indicated. All recordings were performed using the FDSS 7000EX (Hamamatsu) at 37 °C. The FDSS 7000EX is a high-throughput screening (HTS) system for fluorescence and luminescence assays. It enables real-time kinetic measurements of adherent cells in 96- and 384-well plates. The built-in pipettor enables simultaneous compound dispensing across the whole cell-culture plate while recording changes in fluorescence at the same time. For experiments described in this paper the excitation/emission wavelengths were 480/540nm for both GCaMP6f and Calcium6 dye. Both Calcium6 dye (Molecular Devices) and the genetically encoded GCaMP6f are indicators of fluorescence-intensity, which measure changes in calcium concentration by a single fluorescent signal. Cocultures were imaged at 37 °C in calcium imaging buffer (25 mM HEPES, 150 mM NaCl, 8 mM KCl, 1 mM MgCl_2_, 10 mM glucose, 4 mM CaCl_2_, pH to 7.2–7.4) as follows: GCaMP6f fluorescence was recorded for 5 or 10 min at a frame rate of 5 frames/s with binning 2x2 for standard recording and at a frame rate of 8 frames/s binning 1x1 for pixel-resolution. Compounds were prepared in calcium imaging buffer containing 0.1% BSA and were added during acquisition, 5 min after recording start. For calcium imaging assays performed with Calcium-dye, cocultures were incubated for 90 min at 37 °C, 5% CO_2_ with Calcium6 dye (Molecular Devices) dissolved in calcium imaging buffer then fluorescence was recorded as described above.

### 2.7 Calcium imaging data analysis

For the detection of network spikes, the function find_peaks() from the Scipy package (Virtanen et al., 2020) in Python was used and peak prominences of the calcium signals were extracted for each well. Global detection thresholds for each plate were set based on the background signal from empty wells at the outer edges of each plate. From the total number of network spikes in a given time period the frequency was calculated. For analysis of calcium traces at pixel-resolution, Scipy find_peaks() function was used. A prominence-based detection threshold of 0.4 was applied for local field spike detection, unless otherwise mentioned.

### 2.8 Immune-cytochemistry

After removal of medium or calcium imaging buffer, the cocultures were fixed with 4% PFA for 10 min at room temperature (RT), then washed three times with 1xPBS. Blocking and permeabilization were done by 1h incubation at RT with 1x PBS containing 0.3% Triton X-100 and 5% Goat serum. Primary antibodies were diluted in blocking buffer (1% BSA and 0.3% Triton X-100 in PBS) at 4 °C for 16-20h then washed 3 times with 1xPBS. Secondary antibodies incubation was done for 1h at RT in blocking buffer containing Hoechst 33342 followed by 4 washes in 1× PBS. Fluorescence acquisition was done using the Yokogawa CQ1 confocal imaging system. Antibodies are listed in Table S1.

### 2.9 Nuclei counting

As part of the antibody-staining protocol above the plates were stained with Hoechst. Images were acquired on the Yokogawa CQ1 confocoal imaging system with at least 6 planes with 10 µm Z-range of at least 6 fields of view per well). Nuclei were detected in QuPath (0.6.0-SNAPSHOT) with the Stardist extension (Bankhead et al., 2017; Schmidt et al., 2018; Weigert & Schmidt, 2022) and subsequently classified. For initial assessment, the classifier used was trained and validated on images of neuronal and astrocyte nuclei stainings based on RFP (GLUT iN) and Sox9 (astrocyte marker, pA) expression, respectively, and dead cell nuclei (excluded). The accuracy of the method was 98.6 % for neurons, and 87.8% of astrocytes. Once validated, the algorithm was trained on data of each experiment. This included annotation of on average more than 200 nuclear objects across 3-10 images, followed by iterative cycles of the user correcting mislabeling by QuPath and further improve the performance.

### 2.10 Enzyme-linked immunosorbent assay (ELISA) for GABA

GABA levels in the culture medium were measured with the human GABA ELISA kit (ImmusMol) in accordance with the manufacturer’s instructions.

### 2.11 Statistical analysis and graphing

All measurements and comparisons were repeated in multiple independent experiments, as listed in Table S2. Representative experiments are shown. Pooling of data across experiments was purposefully avoided in order to display the variability of data across experiments and wells, which needs consideration given the length and complexity of these multi-week cell culture assays. Endpoint measurements were performed between d28 and d31 of coculture, as mentioned. All comparisons were restricted to within-experiment. Statistical tests were used as indicated and corrected for multiple comparisons (Bonferroni correction). Two-way ANOVA test was performed using the statsmodels.stats.anova.anova_lm() function. Posthoc analyses (Tukey’s multiple comparisons test) of the ANOVA tests were done in Graphpad Prism. Scipy Statistics TTEST_ind() unpaired T-test was applied for across-well comparisons. Scipy Statistics TTEST_rel() paired T-test was applied for within-well comparisons for drug treatment versus baseline activity.

Graphing was done using python seaborn statistical data visualization modules including heatmap(), clustermap(), lineplot(), stripplot(), boxplot(). Box plots denote 2^nd^ and 3^rd^ quartile (semitransparent box area), with bars extending to 1.5 times the interquartile range of the lower and upper quartile (IQR). Individual datapoints are overlaid as dots. Line plots denote the 95% confidence interval around the mean, either as error bars or semi-transparent area, unless otherwise indicated. The sklearn.manifold.TSNE() function was used for t-distributed stochastic neighborhood embedding (t-SNE) (Maaten & Hinton, 2008; Wolf et al., 2018), and t-SNE coordinates transferred to Scanpy.pl.tsne() for overlay with gene expression data. Scipy.stats.entropy() was used to calculate the relative entropy of gene expression distributions. Correlation analyses used the pandas.DataFrame.corr() function.

## 3 Results

### 3.1 Development of an all-human in vitro calcium oscillation assay

To investigate key parameters relevant for calcium signals from in vitro human neural networks we opted for a basic assay setup comprised of excitatory GLUT iN and astrocytes, which was adapted from previous reports (Bassil et al., 2021; Sun & Südhof, 2021; Zhang et al., 2013). Neurons were generated by reprogramming of a stable iPSC line (BIONi010-C-13) using inducible overexpression of the human neurogenic transcription factor NGN2, as previously described and characterized including by electrophysiology (Figure 1A)(Schmid et al., 2021; Shih et al., 2021). To enable measurement of calcium signals exclusively from the GLUT iN, we modified these neurogenic iPSC to express a single copy of GCaMP6f genetically encoded calcium indicator coupled to a red fluorescent protein (RFP) from the ROSA26 locus under control of a constitutive CAGGS promoter (Figure 1A, Figure S1A)(Bertero et al., 2016; Chen et al., 2013; Hitoshi et al., 1991). This new iPSC line was termed Bi-GCaMP6f-2A-RFP. RFP expression provided a means for monitoring overall neuronal density and distribution over time of culture in each well. At the end of the assay, between d28 to d31, calcium imaging was performed on an automated plate reader. By this stage, neurons had developed dense neurite networks and robustly expressed GCaMP6f (Figure 1B).

**Figure 1.**
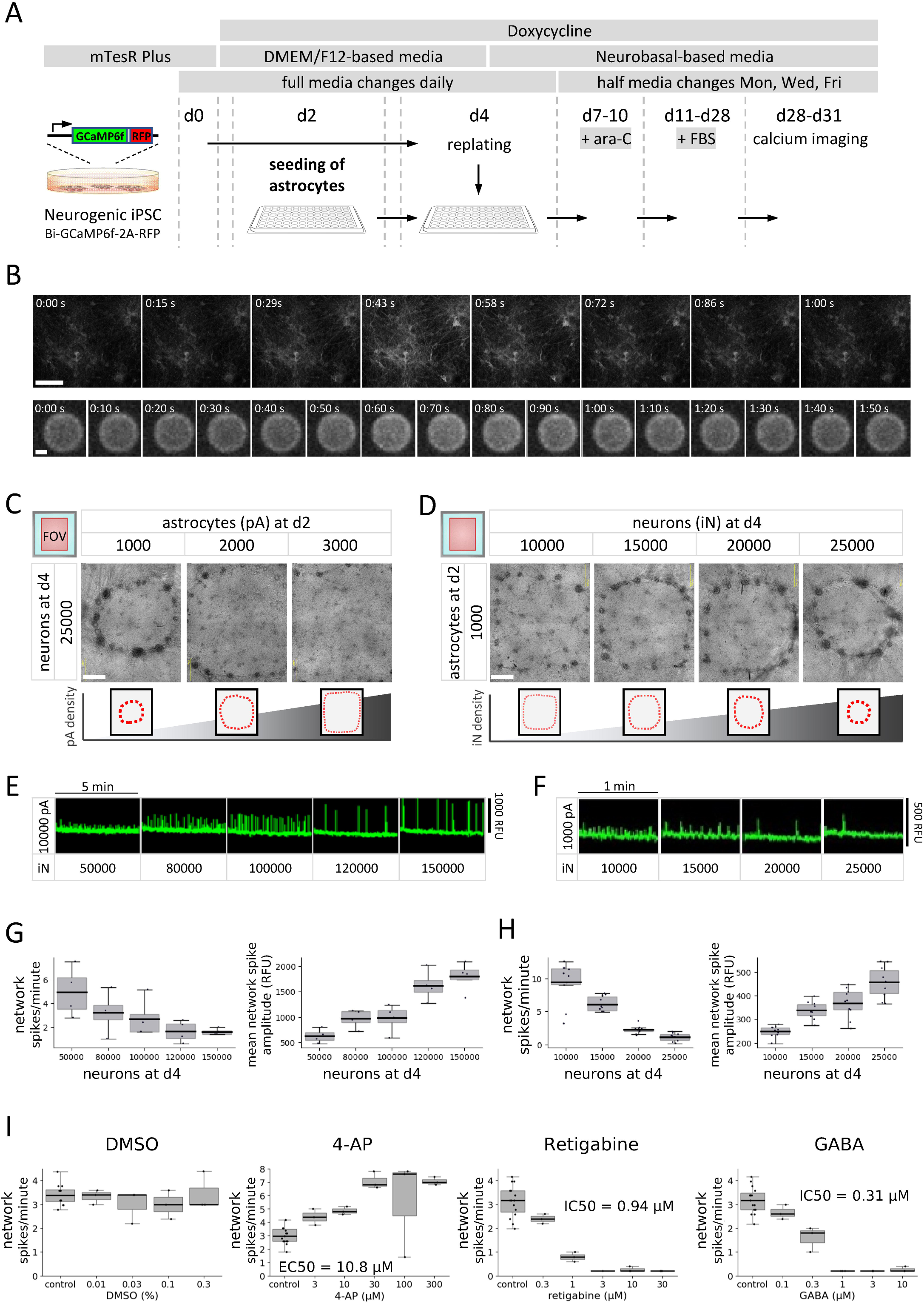
Development of an all-human invitro calcium oscillation assay. A) Schematic overview of calcium assay. Media types, key ingredients, and schedule of media renewal are underlaid gray. Time is denoted as day (d) in culture starting with single-cell seeding of iPSC. B) Examples of GCaMP6f fluorescence across a calcium spike as detected at single-cell resolution on a spinning-disc confocal imaging system (top) and on an automated plate reader (bottom). Scale bar = 100 μm (top) and 2 mm (bottom). (C-D) Circular super-clustering of GLUT iN in 384-well coculture, in dependence of pA seeding density (C), and GLUT iN seeding density (D). Scale bars = 500 μm. Schematic drawings at the bottom illustrate the relationship of cellular distribution and cell type density. (E-F) Excerpt of baseline traces for increasing GLUT iN seeding density at fixed pA density in 96-well (E) and 384-well (F) culture formats visualize the dependence of firing patterns on GLUT iN numbers. Signal correction: SU spatial uniformity. (G-H) Dependence of network spike frequency (left) and mean network spike amplitude (right) on GLUT iN seeding density, at fixed pA seeding densities of 10000 and 1000 per well as in 96-well (G) and 384-well (H) culture formats, respectively. Only wells with fully adherent cells were included in the analysis. N= 4 wells for each neuron density (G) and 10 wells each (H), respectively. I) Concentration-dependent modulation of network activity by potassium channel blocker 4-aminopyridine (4-AP), K_V_7 opener retigabine, and gamma-Aminobutyric Acid (GABA). DMSO served as vehicle. Shown is the network spike frequency (For amplitude see Figure S2). Boxplots denote 2^nd^ and 3^rd^ quartile (gray), with bars extending to 1.5 times of the IQR. Individual datapoints are overlaid as dots.

The early phase of NGN2-driven neural reprogramming using the parent neurogenic iPSC line BIONi010-C-13 has previously been described (Schmid et al., 2021). In terms of cell morphology and density between d0 and d4, cells of our modified Bi-GCaMP6f-2A-RFP iPSC line behaved comparably in our experimental setup, displaying dispersion of iPSC colonies within 1-2 days of doxycycline-induced NGN2 over-expression, and a rapid decline of proliferation as well as appearance of neuronal-like cell morphology by d4 (Figure S1B). We compared the coculture phase of neurons with astrocytes (d4 onward) in 96-well and 384-well plate formats. Two widely used coating conditions, laminin versus Matrigel were tested. Due to lower viscosity and associated ease-of-handling, Matrigel was replaced by the similar Geltrex matrix when switching to 384-well plate format. Coculture conditions were tested at different ratios and densities of neurons and astrocytes. Overall, cocultures behaved similarly on the tested coating conditions for various parameters (Figure S1C, D, G). LDH activity remained overall low, except for a spike upon treatment with Ara-C (d7-d10), which declined after gradual washout with subsequent half-media changes, thus indicating an AraC-induced loss of cells (Figure S1C). Furthermore, average LDH activity across time was more strongly impacted by the number of initially seeded astrocytes at d2 (p_ANOVA_ <0.0001 both for Matrigel and laminin) than by the number of replated neurons at d4 (p_ANOVA_= 0.935 and 0.005, on Matrigel and laminin, respectively) without interaction between the two (p_ANOVA_ = 0.105 and 0.994 on Matrigel and laminin, respectively). These results indicate that the major cell type retaining some degree of cell proliferation into week 2 of the assay are the pA and, consistent with this, that neuron viability is comparably less affected by Ara-C treatment.

### 3.2 The cell type ratio determines cellular distribution and surface adherence

Over the course of 3-4 weeks of coculture, the cellular composition (i.e., ratio and seeding densities of GLUT iN and pA) affected the cellular distribution and the attachment to the well surface, as observed by monitoring of the RFP expression in the GLUT iN (Figure S1D). In the 384-well plate format, astrocyte numbers below 2000 per well resulted in redistribution of neural networks to form clusters and further condense into a ring-shaped “super-cluster” that developed over the course of the coculture phase in an outside-in fashion (Figure 1C, D). A similar phenomenon was observed in 96-well plate format at pA densities lower than 10000 (Figure S1E). Monitoring the 384-well cocultures over time by microscopy showed full adherence in all observed wells at low astrocyte numbers of 1000 seeded pA/well and for both coating conditions over the course of 3.5 weeks (10 of 10 replicate wells, Figure S1F, G). In contrast, for cocultures with high astrocyte number up to all 10 replicate wells showed full detachment of cells from the well surface by d31. This effect was most prominent at low neuron-to-astrocyte (iN/pA) ratio (10000 neurons and 3000 astrocytes per well), with 9 and 10 out of 10 replicate wells detached for laminin and Geltrex coating, respectively. Overall, cell adherence was less vulnerable to changes in cellular composition in 96-well compared to 384-well format (not shown). These findings show that cell type composition determines cellular distribution and adherence and thus, may influence neural network development and function. They further underline the importance of careful monitoring of human CNS-cell cocultures, particularly also in an automated environment, and of building an understanding of how unexpected events affect various endpoint measurements. Given that coating conditions showed minor impact on above analyzed culture parameters, and due to ease of handling, we performed subsequent experiments on laminin, unless otherwise mentioned.

### 3.3 The cell type ratio affects network firing patterns

GCaMP6f signal revealed network-wide firing properties of the various neuron-astrocyte coculture conditions. Baseline neural network activity was recorded for 5 min, followed by acute treatment with neuroactive compounds and 5 min recording of the ensuing drug response over time. Recordings were performed in stimulating buffer conditions adapted from Sun and colleagues (Sun & Südhof, 2021) that enable spontaneous network oscillations. Baseline readings showed consistent network activity with various network spike frequencies and mean network spike amplitudes depending on the coculture composition and culture format (Figure 1E, F). Overall, the frequency ranges in 96-well and 384-well plate formats were comparable, as were spike amplitudes, considering the differences in area and numbers of neurons contributing to the signal (Figure 1G, H). Analysis of network activity patterns revealed a major impact of neuron density on network spike frequency and amplitudes showing a negative correlation with the former (Pearson r of -0.72 and -0.64 for 96-well and 384-well plate, respectively) and a positive correlation with the latter (Pearson r of 0.65 and 0.71 for 96-well and 384-well plate, respectively, Figure 1 G, H). A similar impact was seen for total cell density and GLUT iN/pA cell type ratio. For the latter parameters, results may be explained largely by their strong dependence on the GLUT iN numbers, which exceeded the pA numbers considerably. The impact of pA density on network activity was less clear as a positive correlation of spike frequency with pA density found in the 384-well plate format (Pearson r = 0.38) was not detected in the 96-well plate (Pearson r = -0.073). There was a slight negative correlation of mean spike amplitude with pA density in the 96-well plate and in the 384-well plate format (Pearson r = -0.36 and -0.2, respectively). The overall negative correlation of spike frequency and amplitude across various seeded GLUT iN numbers is also highlighted by graphing the two parameters at constant pA density (Figure S2A). We repeated above experiments with additional GLUT iN/astrocyte ratios to further refine the cell type ratios and confirmed optimal seeding ratios of 15000 GLUT iN and 2000 to 2500 pA for 384-well format and 90000 to 100000 GLUT iN to 10000 pA for 96-well format plates (not shown). Further experiments were performed under these conditions, unless otherwise mentioned.

Next, we tested responses of our system to modulators of neuronal network firing (Figure 1I and Figure S2B, C). First, we established that DMSO, a widely used dissolution agent, did neither impact the mean network spike frequency nor amplitude up to 0.3%. We then demonstrated that our neurons have functional voltage gated potassium channels (K_V_). The K_V_ antagonist 4-aminopyridine (4-AP) increased the mean network spike frequency and reduced the network spike amplitudes in a dose-dependent manner. In contrast, the KCNQ channel opener retigabine led to a dose-dependent reduction in the mean network spike frequency (IC50 = 0.94 µM), whereas the network spike amplitudes were not significantly changed. GABA, the endogenous ligand for GABA receptors, produced a dose-dependent reduction in spontaneous neural network activity (IC50 = 0.31 µM) with only a small effect on calcium spike amplitudes. Lastly, spontaneous neuronal activity was also looked at in the context of Neurobasal/B27 PLUS medium (Figure S2D). However, this culture condition failed to produce robust synchronized firing and their depolarizing response to GABA suggested an immature state of neural networks around 4 weeks in culture.

These results reveal a sensitivity of neural network firing on neuron astrocyte ratios, while two tested coating conditions did not show differential impact. Furthermore, the typical responses of the GLUT iN in our cocultures with pA to modulators of voltage-dependent potassium channels and to GABA suggest the presence of relatively mature, glutamatergic neuronal networks at the time of calcium measurements.

### 3.4 DAPT negatively impacts astrocyte viability

Previous work used the γ-secretase/Notch-signaling inhibitor N-[N-(3,5-difluorophenacetyl)-L-alanyl]-S-phenylglycine t-butyl ester (DAPT) to promote conversion of precursor cells into postmitotic GLUT and GABA iN (Meijer et al., 2019; Rhee et al., 2019). Furthermore, chronic exposure to DAPT led to altered synaptic function in GLUT iN via changes in cholesterol metabolism, including evidence for enhanced excitatory synapse formation (Essayan-Perez & Südhof, 2023). We therefore wanted to test whether DAPT had a beneficial effect on our culture system with respect to neuronal maturation and/or synapse formation. Given evidence from previous observations and LDH measurements that the pA continued proliferating in the days after seeding and prior to ara-C treatment, the actual cell type ratio post-ara-C treatment was not expected to match the original seeding ratios. Therefore, we needed a method to precisely assess final cell numbers, ideally at cell-type resolution. We developed a method to assess the GLUT iN/pA ratio at the end of experiments by harnessing the different nuclear appearances of neurons and astrocytes (Figure 2A). In our cultures, neuronal nuclei were small, roundish, and stained brightly with DNA-intercalating dyes, while astrocyte nuclei were bigger, of elongated oval shape, and stained dimmer compared to neuronal nuclei (Figure S3A). Exploiting these features, we trained a classification algorithm in Qupath (Bankhead et al., 2017) with the StarDist extension (Schmidt et al., 2018; Weigert & Schmidt, 2022), for automated nuclei detection and quantified cell type ratios at d28, with and without acute 10 µM DAPT treatment at d4, similar to what was previously described. Acute treatment with DAPT at d4 resulted in a significant increase in neuron numbers (mean ±std) by d28 compared to control (998.4 ±196.3 /mm² vs 752.4 ±196.3 /mm², p_TTEST_ <0.0001) (Figure 2B). At 10.6276 mm^2^ well area of a 384-well plate this translated to 71% and 53% GLUT iN survival at d28 compared to 15000 GLUT iN seeded at d4 for DAPT treatment vs control, respectively. In turn, we were surprised to find a significantly lower number of astrocytes upon DAPT treatment compared to control (194.4 ±45.1 /mm² vs 300.3 ±64.4 /mm², p_TTEST_ <0.0001), which translated to 82.6% and 127.6% pA survival at d28, compared to 2500 pA seeded at d2 (Figure 2B). These numbers also led to the GLUT iN/pA cell-type ratios at d28 of 5.3 ±1.5 and 2.6 ±1.0 for DAPT-treated vs control cultures (Figure 2C). Due to total cell density mainly determined by the more abundant neurons, the total survival rates, which we defined as total cell number at the end of the assay compared to total number of seeded cells (pA at d2 and GLUT iN at d4), were at 0.75 ±0.12 and 0.66 ±0.14 for DAPT-treated vs control cultures (Figure 2C). DAPT treatment in this experiment overlapped with remaining proliferative activity of pA between seeding at d2 and ara-C treatment starting at d7. We therefore tested whether the reduced pA numbers in DAPT-treated cultures resulted from interference of DAPT with proliferation or alternatively, from death of pA. Realtime monitoring of cell apoptosis in pA monocultures (caspase 3/7 stain) showed an increase in caspase 3/7 activity and a concomitant reduction of cell confluence (Figure 2D, E). Furthermore, the timeline of the live-monitoring results suggested an immediate impact of DAPT on pA viability, which faded after 24 hours. These findings suggest that, while DAPT may have a positive effect on neuron survival, it negatively affects astrocyte viability in GLUT iN cocultures with pA.

**Figure 2.**
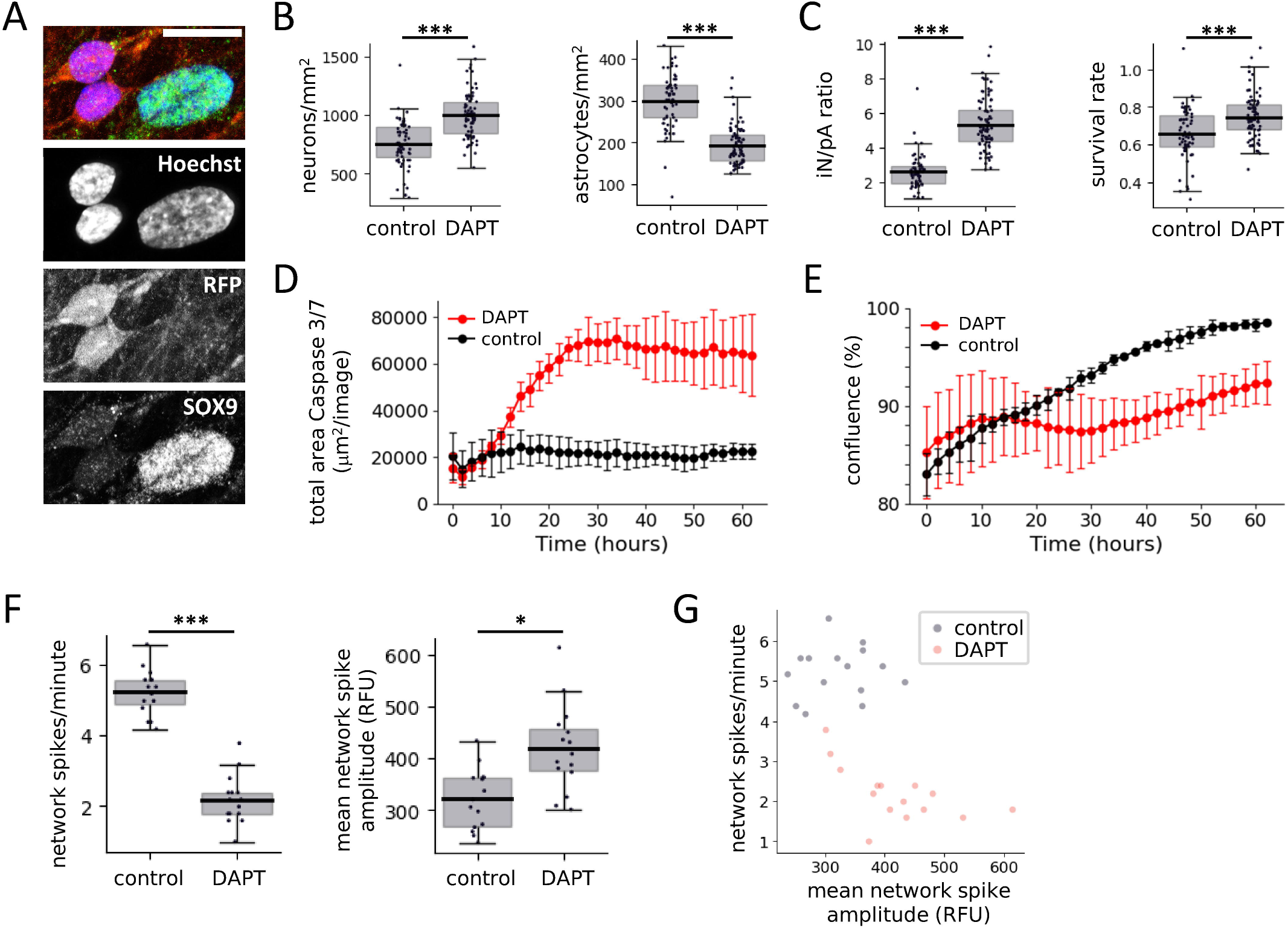
Impact of DAPT on coculture of GLUT iN with pA. A) Fluorescence microscope images illustrate the different nuclear appearance (Hoechst nuclear stain) of iN (identified based on RFP expression) and pA (identified based on expression of astrocyte marker SOX9). Scale bar = 20 μm. B) Quantification of GLUT iN (left) and pA (right) based on cell type-specific nuclear appearance. C) The final iN-to-pA was calculated (left), as well as the survival rate as defined by total cell number at the end of the assay compared to total number of seeded cells. N=80 wells per group. (D-E) Activated Caspase 3/7 area (D) and confluence (E) of pA mono-culture with DAPT treatment and without. N = 3 wells per condition, Error bars denote the 95 % confidence interval. F) Impact of DAPT treatment on neural network activity as measured by GCaMP6f. N= 15 wells per condition. Boxplots denote 2^nd^ and 3^rd^ quartile (gray), with bars extending to 1.5 times of the IQR. Individual datapoints are overlaid as dots. G) Relationship of network spike frequency and mean network spike amplitude shown in F, with and without DAPT treatment *p<0.05, ***p<0.001.

The addition of DAPT for 3 days to the culture medium also affected the network firing patterns. Using both the genetically encoded calcium indicator GCaMP6f as well as the organic calcium dye Calcium-6, we consistently observed a decrease in network spike amplitudes and an increase in network spike frequency (Figure 2F, G and Figure S3B, C). The presence of DAPT significantly reduced the mean network spike frequency from 5.2±0.7 to 2.2±0.7 network spikes/min for GCaMP6f (p_TTEST_ <0.0001) and from 3.2± 0.4 to 0.9±0.3 network spikes/min for Calcium-6 (p_TTEST_ <0.0001). The exposure of our neuron-astrocyte coculture to DAPT resulted in a significant increase in mean network spike amplitude, namely from 322±58 to 419±84 RFU for GCaMP6f (p_TTEST_ =0.026) and from 2664±462 to 4059±341 RFU for Calcium-6 (p_TTEST_ <0.0001). These differences in network firing of DAPT-treated cocultures are largely consistent with what would be expected from variation of GLUT iN/pA ratios, as observed in our previous experiments. Therefore, DAPT impact on network firing is due likely to early loss of pA and increased survival of GLUT iN.

### 3.5 GABAergic inhibitory neurons increase the network spike frequency and reduce the amplitude

Local GABAergic inhibition plays an important role in shaping the functional properties of neural networks across the brain, including via controlling local and global network excitability and synchronization. GABA-mediated inhibition may also contribute to network dysfunction in a variety of CNS disorders including epilepsy, autism, or schizophrenia. We therefore wanted to study the impact of GABA neurons on the in vitro human neural networks described above.

For exploring the effects of GABA-mediated inhibition in the context of the calcium oscillation assay, we engineered a neurogenic iPSC line for reprogramming of GABA iN. We inserted a transgene for inducible expression the neurogenic transcription factors ASCL1 and DLX2, inducible over-expression of which was previously shown to efficiently reprogram iPSC into GABA iN, using lentiviral transduction or within a stable cell line (Rhee et al., 2019; Song et al., 2021; Yang et al., 2017), into a commercially available iPSC line (Figure 3A). Furthermore, the insertion was carried out in the context of testing the usefulness of a more general approach for efficient introduction of any reprogramming code of interest into an iPSC line (Figure S4A-D). For the latter, we equipped the iPSC with a constitutively active rtTA transgene and a tetracycline-responsive Cre recombinase fusion with nuclear EGFP in the ROSA26 and AAVS1 genomic loci, respectively. The transgenes further contained a combination of incompatible Lox sites targeted by Cre to enable recombinase-mediated cassette exchange and efficient insertion of reprogramming codes in place of the Cre-EGFP fusion coding sequence (Figure S4A, E, F)(Baer & Bode, 2001; Kolb, 2001; O’Gorman et al., 1991).

**Figure 3.**
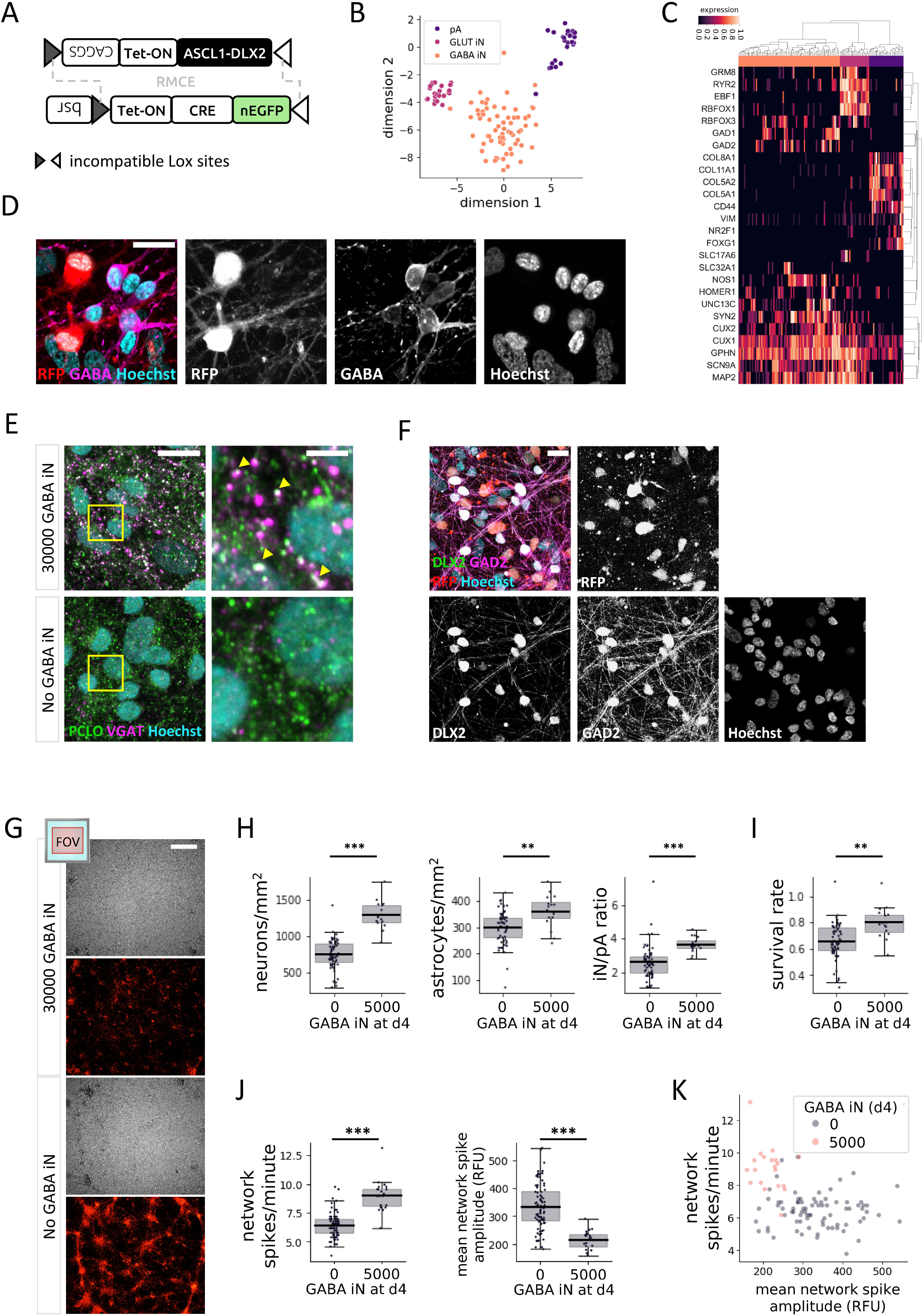
GABA iN affect GLUT iN neural network development and activity. A) Schematic view of genomic landing pad in the AAVS1 locus of the master iPSC line. The Cre recombinase (CRE), connected to nuclear (n) EGFP via a P2A sequence and under control of a tetracycline-sensitive promoter (Tet-ON), is expressed upon treatment with doxycycline. In the presence donor DNA harboring matching incompatible Lox sites, Cre recombinase-mediated cassette exchange (RMCE) inserts the Lox-flanked donor sequence to replace the Cre transgene. Selection is enabled via a CAGGS promoter included in the donor sequence to drive expression of a blasticidin S-resistance coding sequence (*bsr*). (B-C) snRNA-Seq of GABA iN, GLUT iN, and pA from d29 coculture. 2-dimensional T-SNE projection showing the separation of cell types (B), and unsupervised clustering based on cell type-specific marker genes (C). D-F) Immune fluorescence staining of GABAergic marker proteins. D) GABA expression in a coculture of GABA iN, GLUT iN (RFP positive), and pA. Scale bar = 20 μm. E) Expression of piccolo presynaptic cytomatrix protein (PCLO) and vesicular GABA transporter (VGAT) overlaps in presynaptic puncta in cultures with (top, yellow arrowheads), but not in cultures without (bottom) GABA iN. Scale bar = 20 μm (left), and 5 μm on inset magnification (right). F) Expression of DLX2, which is part of the reprogramming code for generation of the GABA iN, overlaps with expression of its target gene glutamic acid decarboxylase GAD2, but not with RFP, which marks the GLUT iN. Scale bar = 20 μm. G) Cellular distribution of 384-well cocultures with 15000 GLUT iN and 2500 pA, in the presence and absence of 5000 GABA iN. Total cells are shown visualized by bright-field. GLUT iN are visualized based on their RFP expression. Field of view (FOV) and area coverage shown on top left. Scale bar = 500 μm. H) Quantification of neurons (combined GLUT and GABA iN, left) and pA (middle) based on cell type-specific nuclear appearance in 384-well cocultures of 15000 GLUT iN and 2500 pA with or without 5000 GABA iN. The final iN/pA ratio is shown on the right. I) The survival rate as defined by total cell number at the end of the assay compared to total number of seeded cells. N=80 wells with GLUT iN and pA only. N=20 wells including 5000 GABA iN per well. J) Calcium imaging of GLUT iN in the presence and absence of GABA iN. Network spike frequency (left) and mean network spike amplitude (right) are shown. K) Scatterplot to visualize the relationship of network spike frequency and mean network spike amplitude in presence and absence of GABA iN. Boxplots denote 2^nd^ and 3^rd^ quartile (gray), with bars extending to 1.5 times of the IQR. Individual datapoints are overlaid as dots. **p<0.01, ***p<0.001.

Following established reprogramming protocols, we generated GABA iN and grew them in coculture on pA, with or without GLUT iN. Four weeks later, we collected cells from 384-well plates to perform multiplexed single-nucleus RNA sequencing (snRNA-seq) using cell multiplexing oligonucleotide (CMO) barcodes to track well origins. For analysis, we pooled data from 2 wells of GABA iN on pA, and 4 wells from coculture also including GLUT iN. CMO labeling efficiency based on 80% confidence was at a relatively low 67%, suggesting that CMOs originally developed for labeling of the plasma membrane may benefit from future adaptations in the context of nuclear membrane properties. In total, contents of 564 droplets were traced back to specific wells of origin. After removal of multiplets and droplets with low signal, visualization in a T-SNE plot distinguished three major cell types matching a total of 124 GABA iN, GLUT iN, and pA (Figure 3B). The distinction is further visualized by clustering based on characteristic marker gene expression (Figure 3C, Figure S4G). Generally, neurons were identified by expression of genes encoding synaptic components (*HOMER1*, *UNC13*, *SYN2*, *GPHN*), cytoskeleton (*MAP2*), and neuron-enriched transcription factors (*RBFOX3*, *CUX1*, *CUX2*). In contrast, the pA transcriptome was enriched for markers of their fetal origin in the forebrain (*FOXG1*), of typical extracellular matrix proteins (genes encoding collagens) and cytoskeletal components (*VIM*), as well as of surface receptors (*CD44*). GLUT and GABA iN differentially expressed surface receptors (*GRM8*, *RYR2*), transcription factors (*EBF1*, *RBFOX1*), and genes playing a role in their respective neurotransmitter metabolism (*GAD1*, *GAD2*) and transport (SLC17A6 versus SLC32A1, respectively). Immune-cytochemistry further confirmed enrichment of GABA in GABA iN versus GLUT iN (Figure 3D). Presynaptic puncta as identified by expression of presynaptic cytomatrix protein piccolo (PCLO) overlapped with staining for vesicular GABA transporter (VGAT/SLC32A1) protein suggesting the presence of GABAergic synaptic terminals in cultures with GABA iN (Figure 3E). Furthermore, the expression of GABAergic neuron marker glutamic acid decarboxylase (GAD2) overlapped with ectopic over-expression of DLX2 as part of the reprogramming code of the GABA iN, and not with RFP, which marked the GLUT iN (Figure 3F). The strong concordance of GAD2 with DLX2 expression is also consistent with studies showing that in rodents, the *Gad2* gene is a direct target of the Dlx2 transcription factor (Le et al., 2017; Pla et al., 2018). We also took this marker analysis as an opportunity to double-check the purity level of the GLUT iN. In the absence of GABA iN, less than 2% of GLUT iN as identified based on RFP expression showed GABA signal, thus confirming the robustness of the NGN2 reprogramming code (Figure S4, H). At the level of other qualitative cell culture properties, the addition of GABA iN reduced the tendency of GLUT iN to form clusters (Figure 3G). In contrast, immune fluorescence staining of pA for S100B and GFAP astrocytic markers did not suggest grossly altered astrocyte morphology or distribution at various GLUT iN densities and with or without GABA iN (Figure S4, I). Nuclei counts at d28 reflected the increased total neuron numbers due to addition of GABA iN, as expected (1294.0 ±195.2 /mm² vs 752.4 ±196.3 /mm², p_TTEST_ <0.0001), and also increased the iN/pA ratio (Figure 3H). However, the degree of increase of total iN in the presence of GABA iN (+72% compared to control) was higher than expected based on the seeding ratios (total 20000 vs 15000 neurons per well or +33.3%). Along the same lines, we observed a significantly higher density of pA in the presence of GABA iN (360.6 ±64.2 /mm² vs 300.3 ±64.4 /mm², p = 0.008), a plus of 20% compared to control. These higher-than-expected numbers were further reflected in a higher survival rate in the presence of GABA iN (0.80 ±0.12), compared to control (0.66 ±0.14) cocultures (Figure 3I). Overall, the apparent positive effect of GABA iN on cell numbers for both neurons and pA, led to final iN/pA ratios of 3.6 vs 2.6 in presence vs absence of GABA iN. This 38% increase roughly matched again the expected 33.3% increase based on the initial number of GABA iN seeded. These results suggest that addition of GABA iN to the coculture of GLUT iN and pA has an overall positive effect on cellular distribution and viability.

The addition of GABA iN resulted in an increase in network spike frequency (no GABA iN vs 5000 GABA iN: 6.4 ±1 network spikes/min vs 9.0 ±1.4 network spikes/min, p_TTEST_ <0.0001) and a decrease in network spike amplitudes (0k GABA IN vs 5000 GABA iN: 335 ±82 RFU vs 215 ±35 RFU, p_TTEST_ <0.0001) compared to cocultures without GABA iN (Figure 3J, K).

### 3.6 Altered neural network activity and reduced synchronization of local field activity in the presence of GABA iN

The increase in GLUT iN network spike frequency observed in the presence of GABA iN, despite having been reported in a similar assay based on fetal rat CNS cells (Xing et al., 2021), is not easily reconciled with GABA-mediated inhibition usually decreasing neuronal excitability. The previous study in rat neuron cultures proposed that GABA-mediated increase in network firing frequency involves reduced synchronization of local firing activity. The resulting decrease of participation of local firing in network spikes could therefore also explain the decreased network spike amplitude we had observed in the presence of GABA iN in our human in vitro system (Figure 3J, K). To further explore this and to assess the observed calcium signals at subnetwork-level, we took to analyzing our plate-reader data at pixel-resolution. This approach turned each pixel into a sensor for local field calcium activity, similar to electrodes in a multi-electrode array (MEA) measuring local field activity based on extracellular voltage-changes. Cocultures of 90000 GLUT iN and 10000 pA, with and without 30000 or 60000 GABA iN were grown in a 96-well format. After recording calcium signals at d29, the gray-scale movies from the plate reader were overlaid with a grid-mask covering a slightly larger area than the actual growth area of the wells, due to angular distortion arising from the camera perspective (Figure S5A). The growth area was estimated based on the autofluorescence signal from dry border wells. Clustering of pixel brightness thus allowed to define growth area, a border zone, and an outside area (Figure S5B). A conservative estimate of the growth area excluding the well border area yielded a coverage of about 400 pixels per well (Figure S5C). This number yields an upper estimated area coverage per pixel of 80425 μm^2^ (∼284 x 284 μm) and, integrated with the mean cell type-specific nuclei counts from this coculture of GLUT iN (1307 ±123) with pA (331 ±21), a mean of 105 GLUT iN and 25 pA covered per pixel.

Analysis of the mean maximum fluorescence intensity across all spikes detected for each pixel of a well enabled identification of pixels with underlying spiking calcium activity. Plotting this signal across wells by pixel rank revealed a drop-off (shoulder) close to 400 pixels, suggesting that most of the pixels within the above estimated growth area of about 400 pixels per well covered at least some neurons that showed calcium spikes. Based on this observation, we focused subsequent analyses on the 400 “most active” pixels, similar to focusing MEA data analysis on “active” electrodes. Heat maps of the top 400 pixels reflected the network spike pattern measured on average across the well (Figure 4A). In addition, they revealed a multitude of un-coordinated local field spikes for each pixel’s location. For ease of reading, the results of the subsequent calcium spike analyses, including mean, standard deviation, fold difference, and statistics shown in the relevant panels of Figure 4 and Figure S5, are detailed in table format (Table 1). The presence of GABA iN in the coculture increased the frequency of network spikes reaching significance for 30000, and significantly reduced their amplitude with 30000 and 60000 GABA iN (Figure S5D, E), largely consistent with experiments in 384-well plate format (compare to Figure 3K). However, this pattern was not followed through at the level of (pixel) spikes reflecting local field activity (Figure 4B). The mean spike frequency of all pixels of a well (averaged across N = 5 wells) was unchanged by the presence of 30000 GABA iN and 60000 GABA iN (Figure 4B, and for graphs visualizing the results per pixel see Figure S5F). The mean amplitude of local spikes averaged across wells showed a significant reduction in the presence of 30000 and 60000 GABA iN (no difference between 30000 and 60000 GABA iN). However, the magnitude of amplitude change of local spikes was far from the magnitude of change observed for network spikes (compare Figure 4B, middle with Figure S5E, right). For example, the amplitude for 30000 GABA iN vs no GABA iN showed a 0.66-fold difference in the case of network spikes, but only a 0.96-fold difference in the case of (local) spikes (Table 1). For local spikes, we also measured the full width at half maximum (FWHM, the spike width at half height), which was reduced in the presence of GABA iN, reaching significance with 60000 GABA iN (including significant difference between 30000 and 60000 GABA iN). The latter could reflect a shortened burst length of local field activity.

**Figure 4.**
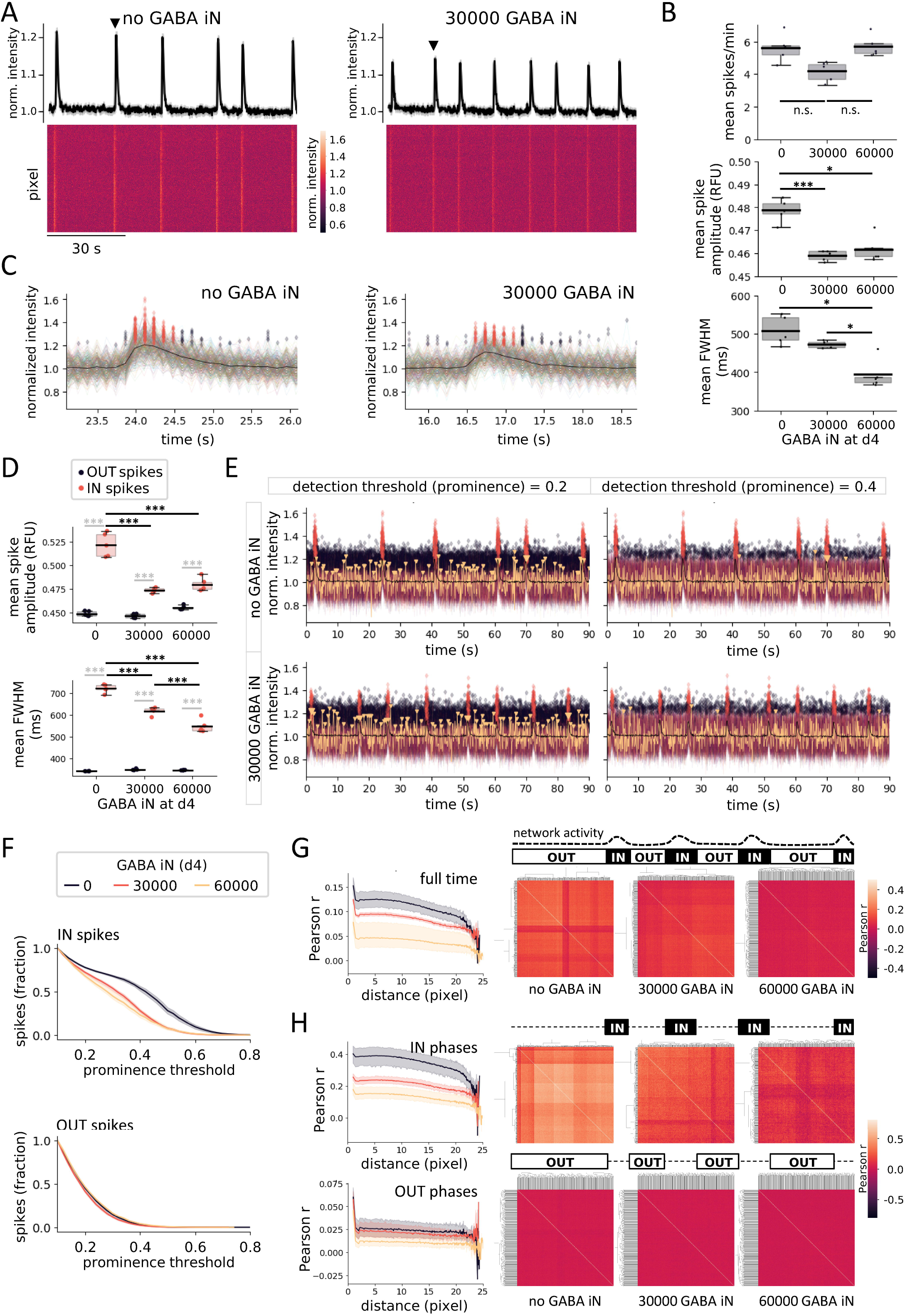
GABA iN impact coincides with GLUT iN network spikes. A) Turning well-based plate reader data into pixel array data for analysis of local field activity in 96-well cocultures of 90000 GLUT iN and 10000 pA, without (left) or with (right) 30000 GABA iN at d29: Normalized GCaMP6f fluorescence intensity heatmaps of the top 400 active pixels over time (bottom). Traces of the average signal reveals the network spikes (top). B) Local field activity: Spike frequency (top), mean spike amplitude (middle), and mean full width at half maximum (FWHM, bottom). N = 5 wells, with 400 pixels averaged per well. For pixel-based graphs, see Figure S5F. C) Pixel-resolution calcium activity around single network spikes (marked with black arrowhead in (A)), in absence (left) and presence (right) of GABA iN. Traces of all top 400 active pixels are overlaid. Black line: mean signal across pixels, transparent red diamonds: Local area (pixel) spikes coinciding with a network spike (IN spikes), black transparent diamonds: spikes that fall outside of network spikes (OUT spikes). D) Mean spike amplitudes (top) and full width at half maximum FWHM (bottom), as separately analyzed for IN spikes and OUT spikes in the presence or absence of GABA iN. E) Visualization of spikes and their coincidence with network spikes at two different detection thresholds, 0.2 (left) and 0.4 (right). For illustration purposes, the calcium trace of a single pixel is shown in yellow with yellow arrowheads denoting IN (large arrowhead) and OUT (small arrowhead) spikes. F) Spike detection in dependence of threshold and shown as fraction of total defined at a low threshold of 0.1 (close to noise, as defined in Figure S5H-I), for IN spikes (top) and OUT spikes (bottom). G) Dependence of pair-wise pixel activity on distance as a measure for spreading and synchronization of local field activity within the well (left). Clustered heatmaps of Pearson correlation coefficients for pair-wise correlation of pixel activity (right). H) Data from (G) separately shown across recording phases coinciding with network spikes (IN phases, top), or not (OUT phases, bottom). Compare schematic illustration above correlation heatmaps with schematic illustration in (G) for definition of recording phases based on network activity. Box plots denote 2^nd^ and 3^rd^ quartile (gray), with bars extending to 1.5 times of the IQR. Individual datapoints are overlaid as dots. Line plots show the 95% confidence interval around the mean as semitransparent area. *p<0.05, **p<0.01, ***p<0.001, n.s. = non-significant.

**Table 1.** Altered network activity and local field activity of GLUT iN in the presence of GABA iN. The table details the mean, standard deviation (std), fold difference, and statistics for frequency, amplitude and FWHM in the presence and absence of GABA iN. Controls are GLUT iN cocultures with pA in the absence of GABA iN. Bold numbers highlight significant changes versus non-significant (n.s.).

| frequency<br>(spikes/min) | well-resolution<br>(network activity) |  |
| --- | --- | --- |
|  | network spikes |  |
|  | mean | std |
| control (no GABA iN) | 3.3 | +/-0.44 |
| 30000 GABA iN | 5.4 | +/-0.69 |
| 60000 GABA iN | 4.3 | +/-1.53 |
| comparisons to control | fold difference | p<sub>TTEST</sub> |
| 30000 GABA iN | 1.64 | 0.004 |
| 60000 GABA iN | 1.30 | n.s. |
| amplitude (RFU) | network spikes |  |
|  | mean | std |
| control | 1598.6 | +/-51.67 |
| 30000 GABA iN | 1048.2 | +/-41.50 |
| 60000 GABA iN | 853.9 | +/-152.42 |
| comparisons to control | fold difference | p<sub>TTEST</sub> |
| 30000 GABA iN | 0.66 | <0.0001 |
| 60000 GABA iN | 0.53 | <0.0001 |

| frequency<br>(spikes/min) | pixel-resolution<br>(local field activity) |  |
| --- | --- | --- |
|  | all spikes |  |
|  | mean | std |
| control | 5.6 | +/-0.86 |
| 30000 GABA iN | 4.2 | +/-0.62 |
| 60000 GABA iN | 5.7 | +/-0.66 |
| comparisons to control | fold difference | p<sub>TTEST</sub> |
| 30000 GABA iN | 0.75 | n.s. |
| 60000 GABA iN | 1.02 | n.s. |
| amplitude<br>(RFU) | all spikes |  |
|  | mean | std |
| control | 0.48 | +/-0.005 |
| 30000 GABA iN | 0.46 | +/-0.002 |
| 60000 GABA iN | 0.46 | +/-0.006 |
| comparisons to control | fold difference | p<sub>TTEST</sub> |
| 30000 GABA iN | 0.96 | 0.0004 |
| 60000 GABA iN | 0.96 | 0.01 |

|  | pixel-resolution (local field activity) |  |  |  |  |  |
| --- | --- | --- | --- | --- | --- | --- |
| amplitude (RFU) | IN spikes |  | OUT spikes |  | comparison IN vs OUT spikes |  |
|  | mean | std | mean | std | fold difference | posthoc p<sub>ANOVA</sub> |
| control | 0.52 | +/-0.013 | 0.45 | +/-0.003 | 1.16 | <0.0001 |
| 30000 GABA iN | 0.47 | +/-0.003 | 0.45 | +/-0.002 | 1.04 | <0.0001 |
| 60000 GABA iN | 0.48 | +/-0.007 | 0.46 | +/-0.002 | 1.04 | <0.0001 |
| comparisons to control | fold difference | posthoc p<sub>ANOVA</sub> | fold difference | posthoc p<sub>ANOVA</sub> |  |  |
| 30000 GABA iN | 0.90 | <0.0001 | 1.00 | n.s. |  |  |
| 60000 GABA iN | 0.92 | <0.0001 | 1.02 | n.s. |  |  |
| FWHM (ms) | IN spikes |  | OUT spikes |  | comparison IN vs OUT spikes |  |
|  | mean | std | mean | std | fold difference | posthoc p<sub>ANOVA</sub> |
| control | 720.7 | +/-19.45 | 340.6 | +/-0.76 | 2.12 | <0.0001 |
| 30000 GABA iN | 617.4 | +/-16.95 | 347.8 | +/-3.37 | 1.78 | <0.0001 |
| 60000 GABA iN | 546.5 | +/-29.78 | 345.3 | +/-1.03 | 1.58 | <0.0001 |
| comparisons to control | fold difference | posthoc p<sub>ANOVA</sub> | fold difference | posthoc p<sub>ANOVA</sub> |  |  |
| 30000 GABA iN | 0.86 | <0.0001 | 1.02 | n.s. |  |  |
| 60000 GABA iN | 0.76 | <0.0001 | 1.01 | n.s. |  |  |

A main difference between the standard, well-resolution data analysis on plate readers and the above proposed pixel-resolution analysis is that the former ignores non-synchronized local field activity, because the consideration of signal intensity across the entire well renders these locally isolated spikes invisible in comparison to well-wide synchronized spikes. In contrast, pixel-resolution analysis treats synchronized and non-synchronized spiking alike (Figure 4C). We therefore wondered whether apparent differences in the magnitude of changes, particularly of the amplitude, seen between network spikes and local spikes was due to the fact that pixel-resolution analysis also includes spikes that are not synchronized, and thus do not contribute to the network spikes as measured at well-resolution. To investigate whether there is a potential difference of GABA iN-mediated changes to local field activity depending on whether such local field activity coincides with network activity or not, we divided the spikes of each pixel into those participating in a network spike (further termed IN spikes), and those that fell outside network spikes (OUT spikes) (Figure 4 C). The mean spike amplitude across wells was significantly higher for IN spikes than compared to OUT spikes (p_ANOVA_ <0.0001) (Figure 4D and Figure S5G). And interestingly, GABA iN only reduced the mean spike amplitude of IN spikes, but not OUT spikes (p_ANOVA_ <0.0001), resulting in a significant interaction of spike type and coculture type (interaction p <0.0001). The FWHM was higher for IN spikes compared to OUT spikes (p_ANOVA_ <0.0001) and selectively reduced by the presence of GABA iN (p_ANOVA_ <0.0001), with significant interaction (interaction p <0.0001). The decreased FWHM for the IN spikes in presence of GABA iN is in line with the observation by (Mossink et al., 2022), who show similar effects for network burst duration using MEA recording, thus demonstrating the power of our imaging-based method despite comparably lower temporal resolution.

Given that spike detection is dependent on amplitude and prominence threshold applied, we also quantified the fraction of spikes detected across thresholds in order to determine threshold-dependent loss of IN spike versus OUT spikes in the presence or absence of GABA IN (Figure 4E, F). We defined a prominence threshold of 0.1, which is just outside of high-intensity noise as defined above (Figure S5H, I), as 100% of spikes detected. This analysis revealed that with increasing spike detection thresholds, IN spikes were lost earlier in the presence of GABA iN compared to GLUT iN cocultures without GABA iN, while OUT spikes were not affected (Figure 4F). This result is consistent with the above-mentioned decrease of spike amplitude at a fixed detection threshold in the presence of GABA iN, selectively observed for IN spikes.

Finally, we harnessed the pixel-based analysis of calcium signals to take a look at network synchronization. The pair-wise correlation of activity patterns for all pixels was calculated and clustered heatmaps created to visualize the patterns of correlation. The presence of GABA iN reduced the correlation coefficients for pairs of pixels and this effect was observed in a GABA iN density-dependent manner (Figure 4G). Adding the distance between pixels based on each pixel’s coordinates on the plate image further allowed for visualization of synchronization in dependence of distance across the network. This analysis showed that the pair-wise correlation coefficients of pixel activity are lower across the well surface in the presence as compared to absence of GABA iN. Based on the IN spike-specific effects of GABA iN observed for spike amplitude and FWHM, we hypothesized that the observed pair-wise correlation might be driven largely by IN spike activity.

Indeed, re-calculating the pair-wise correlation coefficients across pixels separately for local calcium activity overlapping with network spikes (versus non-overlapping) revealed that the correlation was driven largely by phases of network spikes (Figure 4H). In contrast, calcium activity outside network spikes appeared largely uncoordinated with mean Person r about an order of magnitude lower for cocultures without GABA iN. All together these findings indicate that GABA iN have a major impact on the spike properties of local field activity when it participates in synchronized network activity (IN spikes), as opposed to local field activity not participating in network spikes (OUT spikes). Furthermore, this IN spike-selective effect by GABA iN extends to the synchronization of local field activity across the well area, which may result in decreased participation of local field activity in synchronized network activity.

### 3.7 Effects of competitive GABA A receptor inhibition on GLUT iN network activity

Measurements by ELISA of the GABA concentration revealed that the presence of GABA iN significantly increased GABA levels in the culture medium over the course of the 4-week experiment, but also that there was a substantial amount of GABA detectable in the absence of GABA iN (Figure 5A). This could be due to the presence of pA, as astrocytes are capable of producing GABA synthesis through both GAD-dependent and independent pathways (Kilb & Kirischuk, 2022; Kwak et al., 2020; Le Meur et al., 2012). From the snRNA-seq data, we also reasoned that both iN types as well as pA are likely responsive to GABA based on broad expression of genes encoding GABA A receptor subunits as well as GABA B receptors (Figure 5B and Figure S6). To better understand the interplay of GABA iN and GABA signaling in terms of GLUT iN network activity, we therefore decided to test the effects of GABA A receptor blockade with the competitive GABA A receptor antagonist bicuculline. Calcium recording in cocultures of 90000 GLUT iN, 10000 pA, with and without 30000 GABA iN, showed that 5 μM bicuculline treatment did not change the network spike frequency compared to 0.2% DMSO (vehicle) control (p_ANOVA_ = n.s.), while the presence of 30000 GABA iN resulted in a significant increase of the network spike frequency (p_ANOVA_ <0.0001) (Figure 5C, D and Table 2 for mean, standard deviation, fold difference, and posthoc statistics). The mean spike amplitude was significantly increased by bicuculline treatment compared to vehicle control (p_ANOVA_ = 0.0005), but unchanged by the presence of GABA iN (p_ANOVA_ = 0.0676 n.s.). Furthermore, posthoc analysis (Tukey’s multiple comparison test) revealed that the bicuculline effect on network spike amplitude reached significance only in the absence of GABA iN (posthoc p = 0.0013) (Figure 5D, Table 2). In turn, a significant decrease of network spike amplitude in the bicuculline treated, but not the vehicle treated cultures was detected for the presence versus absence of GABA iN (posthoc p = 0.0386). This result could reflect the potential competition between GABA and bicuculline, as predicted from the bicuculline mechanism of action. These results suggest that while GABA iN have an impact on network spike frequency, GABA A receptor blockade by bicuculline does not. In turn, both GABA iN and bicuculline modulate network spike amplitude in opposite directions.

**Figure 5.**
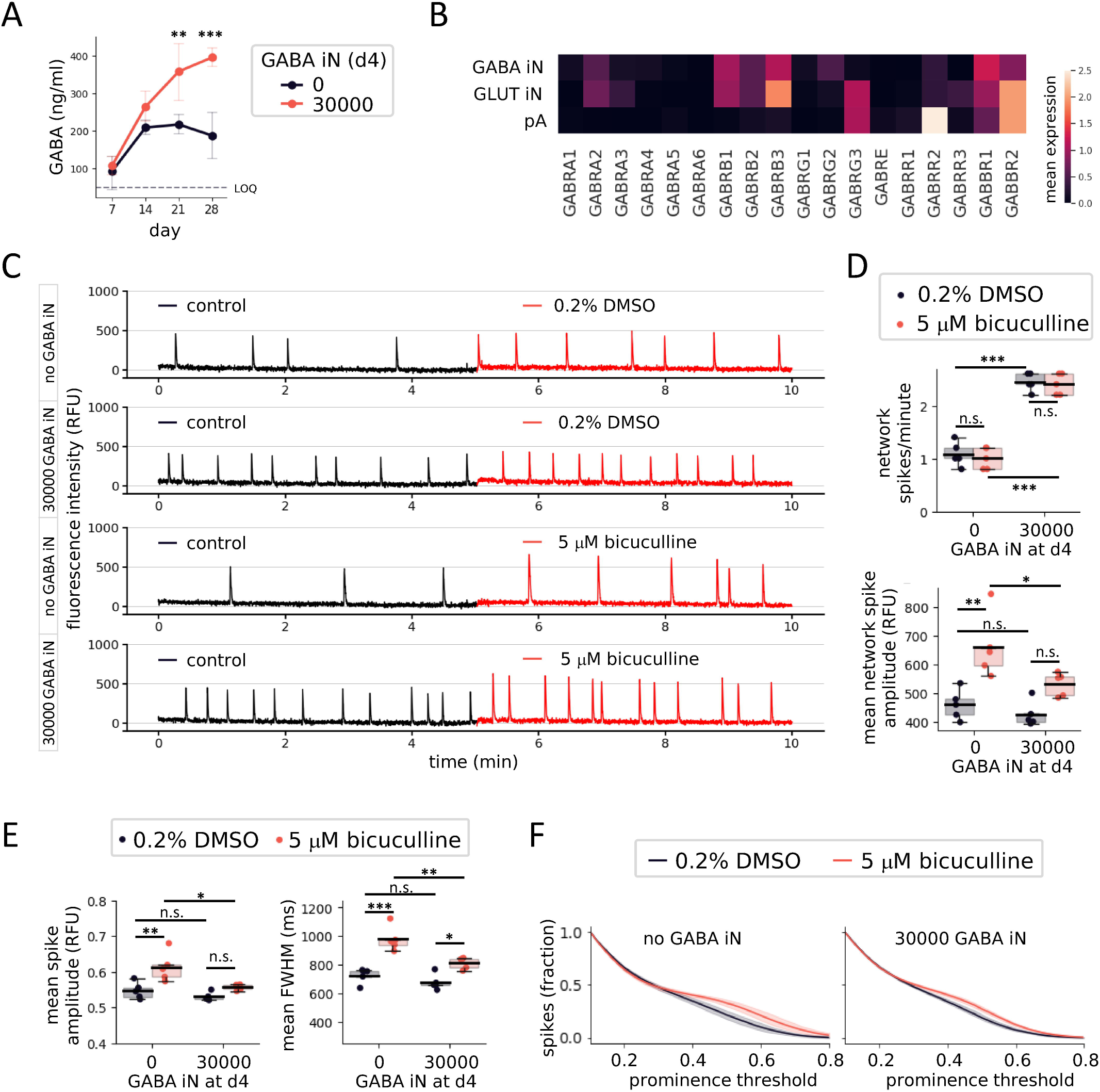
Effects of competitive GABA A receptor inhibition on GLUT iN network activity. D28 coculture of 90000 GLUT iN, 10000 pA, with and without 30000 GABA iN. A) Time course measurements by ELISA of the GABA concentrations. Dashed line denotes the detection limit of the assay. B) Data from snRNA-seq analysis described in Figure 3. The heatmap shows the mean expression (pseudo-bulk analysis) per cell type of genes encoding various GABA A receptor subunits and GABA B receptors. C) Representative calcium traces for wells treated with 5 μM bicuculline or 0.2% DMSO (vehicle). D) Network spike frequency (top) and mean network spike amplitude (bottom) compared between conditions of treatment and GABA iN. E) IN spike amplitude (left) and full width at half maximum (FWHM, right). F) Detection of IN spikes in dependence of threshold and shown as fraction of total defined at a low prominence threshold of 0.1 (close to noise, as defined in Figure S5H-I) The fraction of spikes was calculated for increasing prominence thresholds at a resolution of 0.01 threshold steps. N = 5 wells per condition, Box plots denote 2^nd^ and 3^rd^ quartile (gray), with bars extending to 1.5 times of the IQR. Individual datapoints are overlaid as dots. Line plots show the 95% confidence interval around the mean as semitransparent area. *p<0.05, **p<0.01, ***p<0.001, n.s. = non-significant.

**Table 2.**
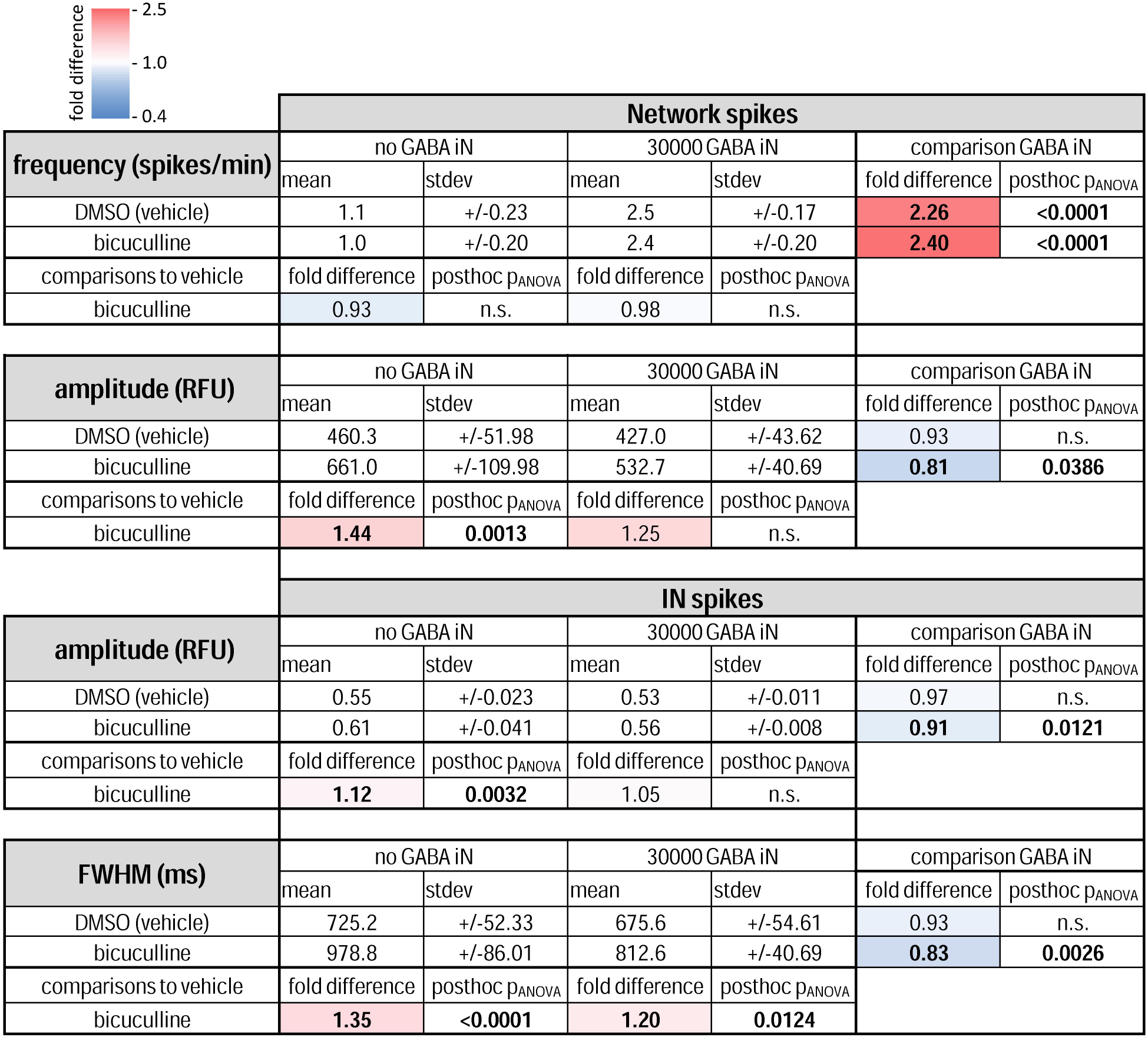
Effects of competitive GABA A receptor inhibition on GLUT iN network activity. The table details the mean, standard deviation (std), fold difference, and statistics for frequency, amplitude and FWHM for bicuculline treatment versus vehicle (DMSO) control in the presence and absence of GABA iN. Bold numbers highlight significant changes versus non-significant (n.s.).

For local field activity measured at pixel-resolution, bicuculline treatment and GABA iN affected exclusively IN spikes, and not OUT spikes (Figure 5E, F, and Figure S7A, B). For IN spikes, the mean spike amplitude was significantly increased by bicuculline (p_ANOVA_ = 0.0065) and decreased in the presence of 30000 GABA iN (p_ANOVA_ = 0.0467), without interaction (interaction p = 0.9460 n.s.) (Figure 5E, Table 2). Similar to the network spike amplitudes, posthoc analyses for IN spike amplitudes showed a significant effect of bicuculline in the absence of GABA iN (posthoc p = 0.0032). And also, a significant decrease of network spike amplitude in the bicuculline treated, but not the vehicle treated cultures was detected for the presence versus absence of GABA iN (posthoc p = 0.0386). The IN spike mean FWHM was significantly increased by bicuculline as compared to vehicle treatment (p_ANOVA_ <0.0001), and significantly increased by the presence vs absence of GABA iN (p_ANOVA_ = 0.0110), without interaction (interaction p = 0.4753 n.s.). Posthoc analysis confirmed the significance of the positive effect of bicuculline treatment on the FWHM, both in cases of presence and absence of GABA iN (posthoc p = 0.0124 and <0.0001). However, the FWHM under bicuculline treatment was significantly smaller in the presence as compared to absence of GABA iN (posthoc p = 0.0026). Also, the quantification of the fraction of spikes detected across thresholds showed that with increasing spike detection thresholds, IN spikes were lost later under bicuculline treatment as compared to vehicle control. A pixel correlation analysis did not yield conclusive results as to the effect of bicuculline (Figure S7C, D) in this experiment. Overall, results suggest an IN spike-specific effect of bicuculline on spike frequency and FWHM. With regards to spike amplitude the findings are largely consistent with the analysis at network spike level suggesting an overall increase of spike amplitudes by bicuculline and a decrease of such effect in the presence of GABA iN. Again, neither bicuculline nor GABA iN showed an impact on OUT spikes (Figure S7A, B).

## 4 Discussion

This study set out to develop a calcium oscillation assay based on coculture of human excitatory neurons and astrocytes, towards scalable applications suitable for high-throughput compound screening. As cellular model for excitatory neurons we chose NGN2-reprogrammed GLUT iN derived from human iPSC. Primary fetal astrocytes, pA, provided glial support for neural network development. The assay was optimized to be compatible with regular weekday working schedules and automated liquid handling.

Our data show that the ratio of astrocytes and neurons affects cell adhesion and distribution on common HTS-compatible optical plates and furthermore, that they shape neural network activity as measured by calcium entry into the neurons. The GLUT iN in our cocultures with pA display a typical response pattern to modulators of key neuronal ion channels including voltage-dependent potassium channels, as well as GABA receptors, suggesting a degree of maturity to serve as functional in vitro models of human excitatory neuron types. The assessment of DAPT, a γ-secretase/Notch-signaling inhibitor, for previously reported promotion of cell cycle exit and maturation of GLUT and GABA iN as well as modulation of synapse formation, was interfered by the compound’s negative impact on astrocyte viability, thus altering the cell type ratios of the coculture system. In this context and to properly compare seeded ratios of astrocytes and neurons at d2 and d4, respectively, with their ratios at the time of functional measurements, an image-based quantification method was developed that takes advantage of the different appearances of astrocytic and neuronal nuclei when stained with nuclear dyes. To include GABAergic neurons in our assay system, we developed a neurogenic iPSC line based on the previously established GABAergic neuron reprogramming code ASCL1+DLX2. Reprogramming of this iPSC line generated GABA iN that express GABAergic neuron markers and increase the GABA concentration in the cell culture medium. Cocultures including these GABA iN displayed an increased network spike frequency and a reduced amplitude compared to cocultures consisting of GLUT iN and pA only. We increased the resolution of the calcium measurements from the plate reader to the level of local field calcium activity by separately analyzing the fluorescence signal of each pixel. This approach revealed local activity outside network spikes (OUT spikes) that are lost in the standard, well-averaged plate reader output. This method revealed that the effect of GABA iN was selective to local field activity overlapping with network spikes (the IN spikes) and that no effect was detected for the temporally independent OUT spikes. Application of the competitive GABA A receptor antagonist did not alter the network spike frequency, but increased the network spike and IN spike amplitudes, while not affecting OUT spike activity.

While it is generally agreed that the formation of functional human neural networks in vitro depends on signals from astrocytes to promote synapse formation and maturation, examples of scalable drug discovery assays based on such human neural networks are still sparse (Bassil et al., 2021; Patzke et al., 2015; Saavedra et al., 2021; Shan et al., 2024). Our work shows that development of such scalable assays is complicated by several factors, including timelines for maturation of neurons, development of neural networks, continued astrocytic support, uncertainties about the cell type ratios in a two-dimensional culture setup, and an incomplete mechanistic understanding of how cellular activity generates synchronized activity of an entire neural network. Our work emphasizes the importance of careful assessment of the ratio of neurons and astrocytes, not just when seeding the cells, but also at the time of functional endpoint measurements. It further adds to the growing toolbox for precise quantification of neurons and astrocytes in such assays by means of differentiating these two cell types based on their nuclear appearance after fluorescent staining. In principle, our nuclei-based cell type quantification is applicable both to fixed and live-stained cultures, thus enabling real-time monitoring of neuron-to-astrocyte ratios in the future. Our study identified a relationship between cell type densities and network firing patterns. In first approximation, the well average calcium signal amplitude may provide an indication of the number of cells contributing to the signal. We further observed concomitant architectural changes including a differential tendency of GLUT iN to form clusters or disperse more evenly over the course of the assay. Therefore, a generalization of a link between network firing and cell type densities may require additional work to better understand how the latter influence network architecture, connectivity, or synapse formation.

Inclusion in the assay protocol of factors that are commonly thought to promote neuronal network maturation, such as DAPT, may need careful testing for their effects on other cell types. Here we show that even a 24-hour exposure to DAPT has dramatic consequences for astrocytic support due to a negative effect specifically on the viability of the pA, in contrast to GLUT iN. The Notch signaling pathway is an important regulator of cell proliferation and maturation. Therefore, while an impact on remaining proliferative activity of astrocytes might have been expected by pharmacological modulation of Notch signaling, it came as a surprise to us that its major impact was on cell viability. The herein reported negative effect of DAPT may merit further exploration, particularly if the compound’s positive effects on neuronal viability and maturation want to be exploited in such human in vitro calcium oscillation assays.

At the network activity level, our data largely showed that GABA iN, somewhat counterintuitively, increase the frequency of network spikes while reducing the network spike amplitude. This is in line with a similar study using cocultures of excitatory and inhibitory neurons (Xing et al., 2021). However, while the electrode readout from MEA has a temporal resolution for isolation of single action potentials from a local field, it is unable to resolve cell type-specific contributions to the observed synchronized network activity. The imaging-based approach we took here, while resolving activity patterns only at the level of action potential bursts, is able to selectively provide signals from the GLUT iN population in the networks, due to cell type-specific expression of the calcium indicator GCaMP6f. Furthermore, we enabled the measurement of local field calcium activity by extracting and analyzing fluorescence intensity data from the plate reader at pixel resolution. This approach enables coverage of local calcium activity with a conservatively estimated 400 pixels per well of a 96-well plate, with the potential for scaling to 384-well format at an estimated coverage of roughly 130 pixels per well for the optical plates used herein (giving or taking of the well boundary area, as described in Figure S5B, which may also differ between round and square shaped wells). These metrics surpass most current MEA formats both in spatial coverage and available well-replicate numbers per plate. Using pixel-resolution data to assess local field calcium activity, we found evidence that GABA iN-mediated effects are largely restricted to time windows covering synchronized network-level neuronal activity, while sparing locally confined calcium spikes that do not coincide with network spikes. The observation that the presence, as compared to absence, of GABA iN came with a steeper detection threshold-dependent loss of IN spikes, and a reduced correlation of local calcium activity across the well may be consistent with findings by Xing and colleagues (Xing et al., 2021) related to reduced network spike participation of single neurons and network desynchronization with increasing inhibitory neuron content in the culture. Furthermore, the decreased FWHM for the IN spikes in presence of GABA iN is in line with the observation by Mossink and colleagues (Mossink et al., 2022), who show similar effects for network burst duration using MEA recording, thus demonstrating the power of our imaging-based method despite comparably lower temporal resolution.

The interpretation of results from treatment with the competitive GABA A receptor antagonist bicuculline is complicated by the fact that (I) even GLUT iN cocultures with pA devoid of GABA iN are not a GABA-free system, as demonstrated by ELISA measurements of GABA, and that (II) all cell types in the co-culture system are likely responsive to GABA, as suggested by gene expression data. Surprisingly, bicuculline had no effect on network spike frequency compared to vehicle treatment, whether or not GABA iN were part of the coculture. Future work should address this phenomenon in more detail. For the network spike and IN spike amplitudes, as well as the IN spike FWHM, bicuculline showed an increase, which was apparently weakened by the presence of GABA iN. Part of an explanation for observed effects at the level of these parameters may be the competition of GABA and bicuculline. The overall opposite effects of GABA iN and GABA A receptor blockade on network spike and IN spike amplitudes, as well as IN spike FWHM and threshold-dependent spike loss, may fit the general idea of GABA shaping these aspects of GLUT iN activity through GABA A receptors. How exactly GABA A receptor blockade changes GLUT iN network activity, either directly or indirectly via modulating GABA iN activity, remains to be determined. The possibility for cell type-specific expression of genetically encoded calcium indicators like GCaMP6f lends itself to future experiments monitoring each cell type’s calcium activity and drug-induced changes thereof.

The analysis of local field activity yielded some interesting observations about IN and OUT spikes that provide an opportunity to speculate about possible differences between the underlying calcium signals and their meaning. OUT spikes generally showed lower variability for spike amplitude and FWHM, compared to IN spikes. In addition, OUT spikes were apparently insensitive to the presence of GABA iN and drug treatment that modulates GABA signaling. Together, these findings could indicate that OUT spikes represent isolated, cell-autonomously generated calcium activity that is not a result of propagating activity from and to neighboring neurons but instead, is driven mainly by the internal state of cells. In such scenario, the IN spike sensitivity to changes in network architecture (e.g. by GABA iN) and to GABA modulation would be explained by their dependence on network activity, synchronization with connected neurons, and propagation of their calcium signals from and to neighboring neurons.

In conclusion, this study contributes to developing functional neural network assays based on human CNS cell types, and to facilitating their application in scalable drug discovery environments. This is of particular relevance in light of initiatives by regulatory agencies, including the FDA Modernization Act 2.0, gaining traction in drug discovery research to increasingly complement and, where possible, replace animal models with human cellular model systems (Zushin et al., 2023). Our work highlights key parameters to be considered when establishing calcium imaging assays to study human neural network function in vitro, and methods to reconcile scaling of optical calcium measurements by microplate reader with a level of resolution enabling the recording of local field activity similar to MEA systems. A major limitation to generalization of this and similar assay protocols remains the primary tissue origin astrocytes, which play a critical role in development and function of neural networks. Anatomical origin as well as preparation inevitably result in batch-specific properties of pA. The replacement of primary cells to build fully iPSC-derived assays of unlimited cellular sourcing should therefore become a primary focus of future research. Protocols for reprogramming iPSC into astrocytes (iA) have been reported and thus offer a promising starting point in this direction (Canals et al., 2018). Fully renewable sourcing of cells is of particular importance for scalability and automation of assays in drug discovery applications. Another point to keep in mind is that for systematic, high-throughput drug testing, short recording periods of calcium signals are of value. However, the benefits of keeping data collection short come at the cost of data points and statistical power, particularly if the neural networks fire at relatively low frequency. For design of drug screening assays, balancing these two aspects therefore needs careful consideration. Finally, to what extent our study, and similar findings by others, have implications for our understanding of neural network activity, synchronization, and GABA action in vivo, including in epilepsy, autism, schizophrenia or other conditions, remains to be seen and may determine the predictive value of human in vitro neuronal model systems for future drug discovery research.

## 5 Conflict of Interest

All authors are or were at the time of the study employees of Idorsia Pharmaceuticals Ltd.

## 6 Author Contributions

Conceptualization (TP)

Data curation (MDH,TP)

Formal analysis (MDH,TP)

Investigation (CD,SS,LL,JM,EG,LT,PB,BR,SB,SW)

Methodology (MDH,TP)

Project administration (TP)

Software (MDH,TP)

Supervision (MDH,EG,PB,TP)

Validation (MDH,TP)

Visualization (MDH,TP)

Writing – original draft (TP)

Writing – review & editing (MDH,CD,SS,LL,JM,EG,LT,PB,BR,SB,SW,TP)

## 7 Acknowledgments

We thank John Fuller and Bruno Sempère for help with the implementation of nuclei quantification by QuPath/StarDist, and Diego Fernandez and John Fuller for help with initial versions of network spike detection using Python.

## 8 Data Availability Statement

The snRNA-seq data are available at the European Nucleotide Archive (ENA) under accession number PRJEB98723.

## Notes

### Competing Interest Statement

The authors have declared no competing interest.

### Summary of Updates

This version of the manuscript has been revised based on peer input. For ease of navigation and readability, results and statistics of Figures 4 and 5 are now summarized in Table format. Figure 5 contain updated experimental data. Corrections were made to typos, and for text to better align with standards in the field.

