## Supplementary Materials for "Development and characterization of a scalable calcium imaging assay using human iPSC-derived neurons"

### *Supplementary Material*

#### 1 Supplementary methods

##### 1.1 Human iPSC culture and engineering

Human iPSCs were cultured in mTeSR Plus on Geltrex coated cell culture vessels (1:100 in DMEM/F12) at 37 °C, 5% $CO_2$ and >90% humidity. Cells were passaged as aggregates by incubating them for 5 min at 37 °C with ReLeSR and seeding them in mTeSR Plus supplemented with 2 µM Thiazovivin for less than 24 h.

For CRISPR-based genome engineering, RNP complexes were prepared in two steps. First, the crRNA and tracrRNA-ATTO 550 were mixed at 5 µM in Neon Resuspension buffer R, heated to 95 °C and slowly cooled down to room temperature. Then the newly formed gRNA was incubated at 4 µM with HiFi Cas9 nuclease V3 at 1.4 µM for 20 min at room temperature. Cells were incubated in mTeSR Plus supplemented with 1x CloneR2 for 1 h, and subsequently detached with Accutase for 8 min at 37 °C. The cells were pelleted at 300 g for 5 min and resuspend in Neon Resuspension buffer R at 60000 cells/µL. 7.5 µl cell suspension was mixed with 5 µL RNP complexes and 2.5 µL donor plasmid (1.5 µg in Neon Resuspension buffer R). 10 µL was aspirated with the Neon pipette and electroporated with a single 30 ms pulse at 1'100 V. Immediately afterwards, the cells were transferred to a Geltrex coated 6-well plate containing mTeSR Plus supplemented with 1x CloneR2 and 20 µM HDR-Enhancer V1 and incubated at 32 °C, 5% $CO_2$. After 2 days, the medium was replaced with mTeSR Plus supplemented with the appropriate selection antibiotic (Puromycin: 0.3 µg/mL / Blasticidin: 2 µg/mL) and the cells were continuously monitored until clonal selection.

For Cre recombinase-mediated cassette exchange (RMCE), cells were treated with 1 µg/mL doxycycline for 24 h before electroporation to induce the expression of Cre. Single-cell suspension in Neon Resuspension buffer R was prepared as described above and mixed with 1.5 µg donor plasmid, diluted in 7.5 µL Neon Resuspension buffer R. 10 µL of the mix was electroporated with a triple 20 ms pulse at 1'200 V. Immediately afterwards, the cells were transferred to a Geltrex coated 6-well plate containing mTeSR Plus supplemented with 1x CloneR2. After 2 days, the medium was replaced with mTeSR Plus supplemented with 2 µg/mL Blasticidin and cells monitored until clonal selection.

Donor DNA constructs were generated by gene synthesis (Azenta). Schematic views are shown for donor constructs for homology-directed repair (HDR) (Fig S1A, Fig S4A,C) and RMCE (Fig 3A). For HDR constructs, the selection cassettes were flanked by compatible Lox sites (LoxFas and LoxN, for the puromycin N-acetyltransferase, *pac*, and blasticidin S deaminase, *bsr*, genes respectively). A Cre recombinase coding sequence was coupled to EGFP via a P2A sequence and put under control of a tetracycline responsive Tet-ON promoter. A reverse Tet transactivator (rtTA) coding sequence was put under control of the constitutive CAGGS promoter (Hitoshi et al., 1991). Selection cassettes flanked by compatible Lox sites were removed in the final master iPSC line by brief exposure to doxycycline to enable their reuse for integration of additional DNA modules. The Cre transgene was flanked by incompatible Lox sites (LoxP and Lox2272). Upstream of the Tet-ON promoter in opposite direction, a *bsr* gene without promoter was placed to enable selection for RMCE upon integration of a reprogramming code. RMCE donor constructs contained the reprogramming code ASCL1+DLX2 (coupled by a 2A sequence) under control of a Tet-ON promoter. Furthermore, a CAGGS promoter was placed upstream in the opposite direction to drive *bsr* and enable selection for genomic integration by RMCE. The donor DNA of the RMCE contructs were flanked by incompatible LoxP and Lox2272 sites matching the replacement cassette in the master iPSC.

For PCR genotyping, cells were washed DPBS and incubated with 80-100 µL/cm$^2$ Lysis solution (Tris-HCl: 10 mM, KCl: 50 mM, $MgCl_2$: 2 mM, Gelatin: 0.1 mg/mL, IGEPAL: 0.45%, Tween-20: 0.45%, Proteinase K: 100 µg/mL) for 3 min. The lysates were collected and incubated at 55 °C for 4 h, followed by Proteinase K deactivation at 95 °C for 10 min. gDNA was then amplified by PCR with Platinum™ Hot Start PCR Master Mix. Reagent item numbers and primer sequences are provided in Table S1.

#### 1.2 Neural reprogramming of human iPSC and neuron-astrocyte co-culture

Human iPSCs were detached with Accutase for 8 min at 37 °C, pelleted at 300 g for 5 min, resuspended in mTeSR Plus supplemented with 2 µM Thiazovivin and seeded on Matrigel coated culture vessels at a density of 10000 cells/cm$^2$. From d1 to d3 the medium was exchanged to reprogramming medium (DMEM/F12 supplemented with N2, NEAA, 10 ng/mL hBDNF, 10 ng/mL hNT-3, 200 ng/mL mouse laminin, and 2 µg/mL doxycycline) and renewed daily. On d4 the young neurons were detached with Accutase for 6 min at 37 °C, pelleted at 300 g for 5 min and either resuspended in Bambanker for cryopreservation or in Neuronal medium (Neurobasal-A supplemented with B27, GlutaMAX, 10 ng/mL hBDNF, 10 ng/mL hNT-3, 200 ng/mL mouse laminin, and 2 µg/mL doxycycline) for co-culture with astrocytes. Reprogramming was monitored by microscopy and samples for viability assays were collected at selected timepoints.

On d2 of reprogramming, human primary astrocytes were seeded on laminin (20 µg/ml in DPBS) coated culture cell culture plates (Perkin Elmer phenoplate) in Astrocyte medium. After 2 days, the medium was switched to Neuronal medium and young neurons (d4) were seeded on top. Half-media changes were performed every 2-3 days. From d7-d10, the neuronal medium was supplemented with 2 µM Ara-C and from d11 onwards with 2.5% FBS. The co-cultures were monitored by microscopy and samples for viability assays were collected at selected timepoints.

#### 1.3 Cell viability assays

LDH assay was performed with the CytoTox 96 non-radioactive assay, according to manufacturer's protocols. Briefly, cell culture conditioned medium was collected and 12.5 µL were transferred in duplicate or in triplicate into a 384 well plate, spun down, and supplemented with 12.5 µL of CytoTox 96 Reagent. After 15-30s mix with a plate shaker, assay plate was incubated 30 minutes at room temperature, light protected. Reaction was stopped by adding 12.5 µL of Stop Solution, mixed 15-30s with a plate shaker, light protected, and absorbance was measured immediately (or within one hour) at 490 nm with the SynergyMX microplate reader, without lid. Cell culture medium served as control. Data processing and analysis were performed according to manufacturer's recommendations.

Incucyte® Caspase-3/7 Green Dye was used according to manufacturer's instructions. Briefly, 5 µM Dye was diluted in culture medium and phase and fluorescent images were acquired with the Incucyte SX5 every 2h. Masks for the cell confluence and the dye fluorescence were directly generated in the Incucyte software.

#### 1.4 Single-Nucleus RNA Sequencing (snRNA-seq) Library Preparation and Sequencing

To prepare single-nuclei suspensions for snRNA-seq, co-cultures plates were placed on ice for 1 min and medium was removed. 20 µL/well of cold Lysis buffer (1x PBS, Tris-HCl 10 mM, NaCl 10 mM, $MgCl_2$ 3mM, NP40 0.025%, pH7.4) were added and cells were kept on ice for 3 min with 3-5 times gently mix using a micropipette. Lysis was stopped by adding 4x lysis buffer volume of cold Nuclei Wash and Resuspension Buffer (NWR Buffer) (1x PBS, 1% BSA, RNase Inhibitor 0.2 U/µL). The nuclei suspensions were collected in a microplate and centrifuged for 10 min at 500 rcf, 4 °C using swinging-bucket rotor. Supernatants were removed and nuclei pellets were resuspended in 180 µL of NWR buffer and then centrifuged at 500 rcf for 10 min at 4 °C. The washing step was repeated once more and the nuclei were resuspended in 1x PBS containing 0.04% BSA.

3' CellPlex Kit Set A was used for nuclei labeling. Before use, Cell Multiplexing Oligo (CMO) were thawed 30 min at room temperature, then vortexed 5 sec at maximum speed and centrifuged briefly for 5 sec. Supernatants were removed and nuclei were resuspended with 20 µL CMO with 15x gentle pipette mix. Labeling was done for 5 min at room temperature. 9x CMO volume of cold NWR Buffer was added and nuclei were centrifuged at 500 rcf for 10 min at 4 °C. Supernatants were completely removed, and nuclei pellets were resuspended in 180 µL of cold NWR Buffer then centrifuged at 500 rcf for 10 min at 4 °C. The washing step was repeated two times for a total of 3 washes. Nuclei concentration was quantified using Luna-FX7 automated cell counter and then diluted to 1500 cells/uL for subsequent snRNA-seq experiment.

##### 1.5 Calcium imaging

Calcium imaging was conducted at d28-31 as indicated. All recordings were performed using the FDSS7000EX (Hamamatsu) at 37 °C. Co-cultures were gently washed with modified Tyrode solution (25 mM HEPES, 150 mM NaCl, 5 mM KCl, 1 mM $MgCl_2$, 10 mM glucose, 2 mM $CaCl_2$, pH 7.2–7.4, pre-warmed to 37 °C), then incubated with calcium imaging buffer (25 mM HEPES, 150 mM NaCl, 8 mM KCl, 1 mM $MgCl_2$, 10 mM glucose, 4 mM $CaCl_2$, pH to 7.2–7.4, pre-warmed to 37 °C). After 20 min equilibration at 37 °C, calcium imaging was performed as follows: GCaMP6f fluorescence was recorded for 5 or 10 min at a frame rate of 5 frames/s with binning 2x2 for standard recording and at a frame rate of 8 frames/s binning 1x1 for pixel-resolution. Compounds were prepared in calcium imaging buffer containing 0.1% BSA and were added during acquisition 5 min after recording start. For calcium imaging assays performed with Calcium-dye, co-cultures were incubated for 90 min at 37 °C, 5% CO2 with Calcium6 dye (Molecular Devices) dissolved in calcium imaging buffer then fluorescence was recorded as described above.

#### 2 Supplementary Figures and Tables

##### 2.1 Supplementary Figures

**Figure S1. Development of an all-human invitro calcium oscillation assay: culture conditions.** A) Genotyping data for the genome-edited ROSA26 locus of the BIONi010-C13 neurogenic iPSC line. Schematic view of DNA construct and primer (blue arrows) alignment sites are shown. Top gel:

5-prime integration site, bottom gel: 3-prime integration site. P: parent cell line BIONi010-C13, W: water control, C: isolated clones (sample including positive and negative). The clone expanded for complete QC and use in experiments is denoted in bold font. For primer sequences, see Table S1 B) Early reprogramming phase (d0-d4) monitored by light microscopy at increasing magnifications. Scale bar = 100 μm. C) Relationship of LDH activity and cell type composition in a 96-well coculture before (d7) and after Ara-C treatment, on laminin (left) and Matrigel (right). Data is normalized on d7 baseline LDH activity. N = 5 wells per condition. Error bars denote the 95% confidence interval around the mean. D) Monitoring of RFP expression over time (top to bottom) for cellular distribution and surface adherence. Blue rectangle marks the determined optimal condition. E) Development of circular super-clustering of GLUT iN over time in 96-well coculture at pA seeding density of 2000/well. (F) Qualitative scoring of cellular adherence. G) Quantification of cellular adherence based on qualitative scoring for 384-well cocultures grown on laminin (left) and Geltrex (right). Scale bars E-F: 500 μm.

**Figure S2. Development of an all-human invitro calcium oscillation assay: calcium imaging.** A) Relationship of network spike frequency and mean network spike amplitude across a range of GLUT iN seeding densities at fixed pA seeding densities of 10000 (top) and 1000 (bottom), in 96-well and 384-well coculture formats, respectively. Only wells with fully adherent cells were included in the analysis. N= 4 wells for each neuron density in 96-well plate wells (top) and N = 10 wells for each neuron density in 384-well plate wells (bottom). B-C) Concentration-response-curves calculated from the data in Figure 1I show that the 384-well plate format is suitable for testing the effects of compounds on spontaneous neuronal network activity in a high-throughput manner (N= 3 wells per compound concentration). D) Example traces from $Ca^{2+}$ imaging show that GABA (10µM, red trace) blocks spontaneous neural network activity (black) of neurons grown in Neurobasal A cell culture medium (left). The neurons grown in Neurobasal Plus medium (right) do not show spontaneous activity at baseline (black), and addition of GABA leads to a single spike resembling glutamate-like depolarization of neural networks.

**Figure S3. Impact of DAPT on coculture of GLUT iN with pA.** A) Representative image of Hoechst nuclear staining of neuron-astrocyte coculture (left) and with mask for nucleus detection and counting using QuPath (right). The green outline denotes astrocytic nuclei, purple outline neuronal nuclei. Scale bar = 20µm. B) Network spike frequency and mean network spike amplitude as measured by Calcium 6 dye. N = 21 wells per condition. C) Relationship of network spike frequency and mean network spike amplitude shown in B, with and without DAPT treatment. Boxplots denote 2$^{nd}$ and 3$^{rd}$ quartile (gray), with bars extending to 1.5 times of the IQR. Individual datapoints are overlaid as dots. ***p<0.001.

**Figure S4. Master iPSC line and recombinase-mediated cassette exchange.** A) Schematic view of the modified *AAVS1* locus in the master iPSC line. The locus enables insertion of any reprogramming code by recombinase-mediated cassette exchange (RMCE) of donor DNA sequences flanked by incompatible Lox sites that match the Lox sites integrated in the genomic landing pad. Cre is expressed upon doxycycline treatment. Removal of the puromycin N-acetyltransferase (*pac*) selection cassette successful modification of the landing pad was enabled by compatible Lox sites targeted and Cre upon doxycycline exposure in the absence of donor DNA. Nuclear (n) RFP under a constitutive promoter was used for proof-of-concept RMCE, replacing nEGFP. Promoters are shown as arrows: Tet-ON (black), PGK (orange), CAGGS (light gray), CAG (dark gray). Primer binding sites are shown as arrowhead in the colors matching subsequent figure panels. B) Genotyping data documenting the correct 5’ and 3’ integration of the genomic landing pad. Labels: iPSC clone (“C”), negative control DNA (“-“), water control (“W”). C) Schematic view (top) and genotyping results (middle and bottom) for a CAGGS promoter-driven reverse Tet transactivator (*rtTA*) transgene to enable doxycycline-induced gene expression from the Tet-ON promoter. Selection for the modified ROSA26 locus was enabled by a blasticidin S-resistance gene (*bsr*) that was flanked by compatible Lox sites to enable removal upon doxycycline-induced expression of Cre recombinase. Genotyping results documenting the correct 5’ (middle) and 3’ (bottom) integration of the genomic landing pad after removal of the *bsr* cassette (dashed outline). Labels: master iPSC clone selected for further experiments (“C”), other positive clones (gray “C”), positive control DNA (“+”), negative control DNA (“-“), water control (“W”). D) Pluripotency marker analysis by immune fluorescence on the master iPSC line. Scale bar = 50 μm. E) Proof-of-concept RMCE: replacement of nEGFP by nRFP by RMCE. Master iPSC were treated with doxycycline for 24 hours before electroporation with the RFP-carrying donor DNA (see also (A) for schematic). Cells were treated for another 24 hours after 7 days of recovery to visualize both the recombined (red) and un-recombined (green) cells. Scale bar = 50 μm. F) DNA gels documenting the insertion of Donor DNA by RMCE. Primer (arrowhead) colors matching schema in (A). Clones indicated by large arrowheads: nRFP proof-of-concept RMCE clone (red), neurogenic iPSC clone used for generation of GABA iN (white). Labels: GABA iN reprogramming code ASCL1+DLX2 (“AD”), parental cell line (“P”), water control (“W”), clones carrying other reprogramming codes (“RC”). G) T-SNE plots overlaid with gene expression showing the expression of markers specific to GLUT iN (top row) and GABA iN (bottom row). Color bar represents log2 normalized expression. H) Immune fluorescence quantification of GABA positive (GABA+) GLUT iN as defined by RFP expression (RFP+). N = 4 wells (12 FOV imaged per well). Images show masks for RFP+ nuclei (middle, green outline) and a rare cell expressing GABA (right, blue outline). Scale bar = 20 μm. H) Morphology of pA as visualized by immune fluorescence staining for the spatially complementary astrocyte markers S100B and GFAP cultured at a density of 10000 cells per well (96-well plate format) in the presence of various GLUT iN densities and without or with 30000 GABA iN. Scale bar = 50 μm.

**Figure S5. Pixel-resolution plate reader measurements for analysis of local field calcium activity.** A) Excerpt of a plate reader image of a 96-well plate overlaid with a mask for automated selection of regions of interest (ROI). ROI cover a bigger area than the actual well bottom to ensure coverage of the entire growth area independent of well position and camera angle (inset). Red arrows indicate corresponding areas as described in (B). B) Clustered heatmap of autofluorescence from a dry (border) well. This visualization enables distinguishing pixels that cover the growth area, the border area of the well, and pixels that are located outside the well. C) Ranked pixel intensities based on the mean maximum spike signal enable identification of pixels with calcium spikes (active pixels). D-E) Standard plate reader data output based on averaged fluorescence intensity for each well. Example traces (D) of cocultures with 90000 GLUT iN and 10000 pA in the absence (left) and presence (right) of 30000 GABA iN. E) Mean network spike frequency (left) and mean network spike amplitudes (right). F-G) Analyses of local field activity shown for each pixel. Visualization of well-averaged data in Figure 4 C by mean pixel average (F, left: mean spike frequency, middle: FWHM, right: mean spike amplitude, N = 2000 pixels, combined from 5 wells), or (G) separately for IN and OUT spikes (G, top: mean spike amplitude, bottom: FWHM). H) Pixel traces for the first 30 seconds of baseline calcium recording to illustrate GCaMP6f signal versus high-intensity autofluorescence signal from dry wells. I) Number of detected spikes in dependence of detection threshold applied. N = 5 wells (400 pixel/ well). Comparison of signals from culture is against the high autofluorescence intensity of dry wells. Box plots denote $2^{nd}$ and $3^{rd}$ quartile (gray), with bars extending to 1.5 times of the IQR. Individual datapoints are overlaid as dots. Line plots show the

95% confidence interval around the mean as semitransparent area. *p<0.05, **p<0.01, ***p<0.001, n.s. = non-significant.

**Figure S6. GABA receptor gene expression in GLUT iN, GABA iN, and pA**. T-SNE plots of snRNA-seq data described in Figure 3. Color heatmap represents log2 normalized expression of genes encoding various GABA A receptor subunits and GABA B receptors. Color bar represents log2 normalized expression.

**Figure S7**. **Effects of competitive GABA A receptor inhibition on GLUT iN network activity**. A) OUT spike amplitude (left) and full width at half maximum (FWHM, right) comparing drug treatment to presence/absence of GABA IN in the coculture. B) Detection of OUT spikes in dependence of threshold and shown as fraction of total defined at a low prominence threshold of 0.1 (close to noise, as defined in Figure S5H-I) The fraction of spikes was calculated for increasing prominence thresholds at a resolution of 0.01 threshold steps. C-D) Pair-wise correlation of pixel intensities over time in dependence of distance between pixels, for OUT spikes (C) and IN spikes (D). N = 5 wells per condition, Box plots denote 2$^{nd}$ and 3$^{rd}$ quartile (gray), with bars extending to 1.5 times of the IQR. Individual datapoints are overlaid as dots. Line plots show the 95% confidence interval around the mean as semitransparent area. *p<0.05, **p<0.01, ***p<0.001, n.s. = non-significant.
