## Supplementary figures and images for "Development and characterization of a scalable calcium imaging assay using human iPSC-derived neurons"

### Figure S1

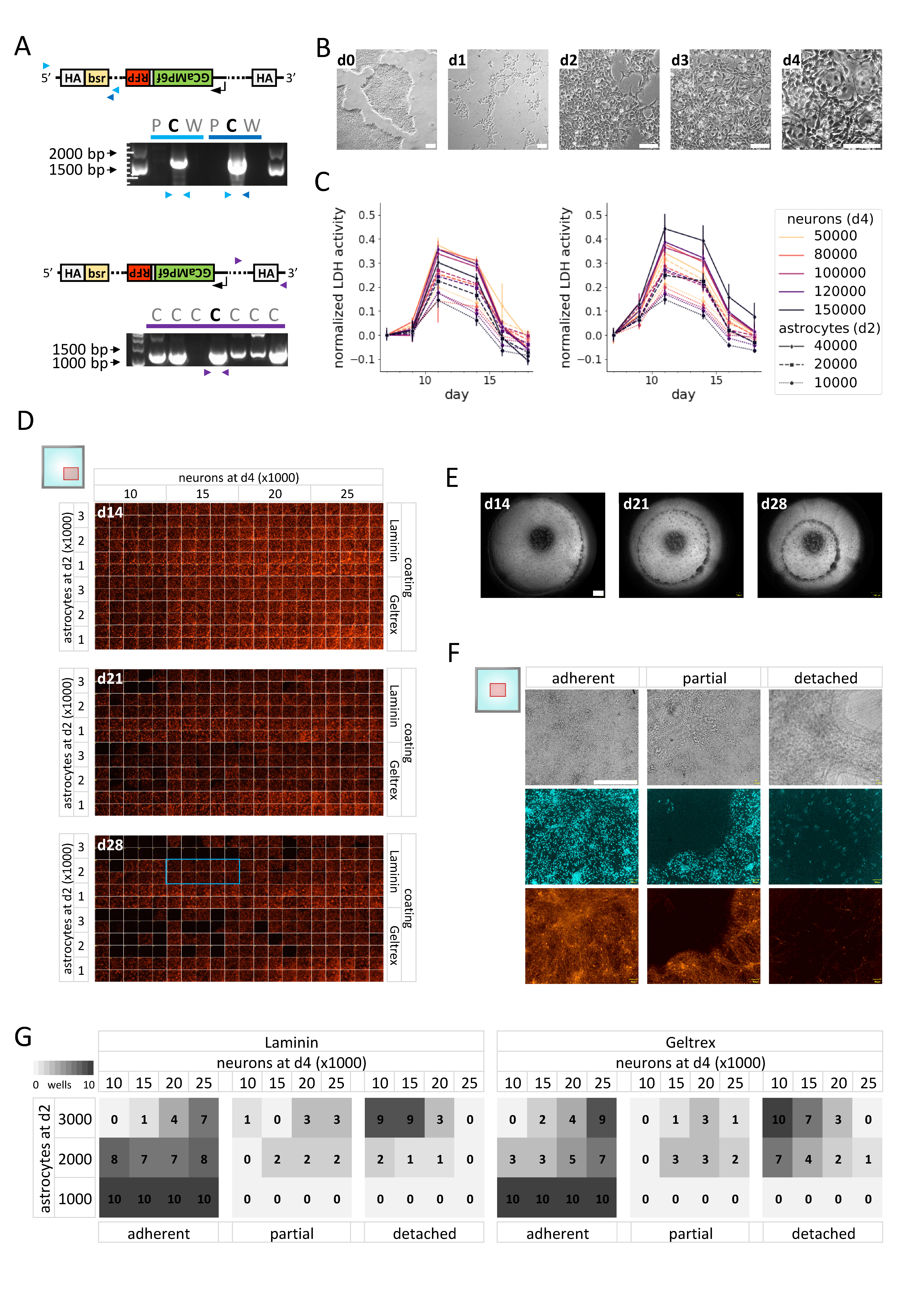

### Figure S2

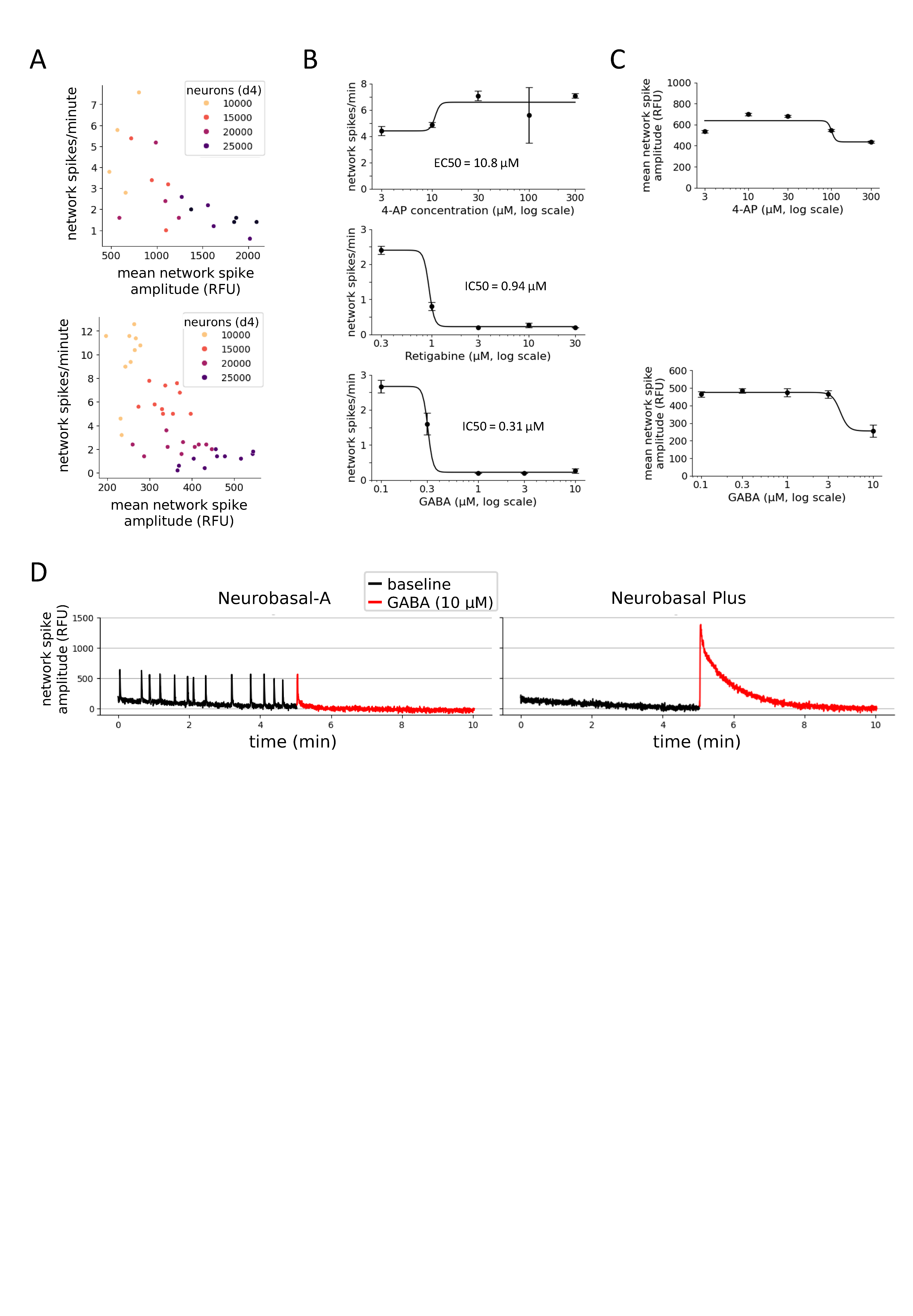

### Figure S3

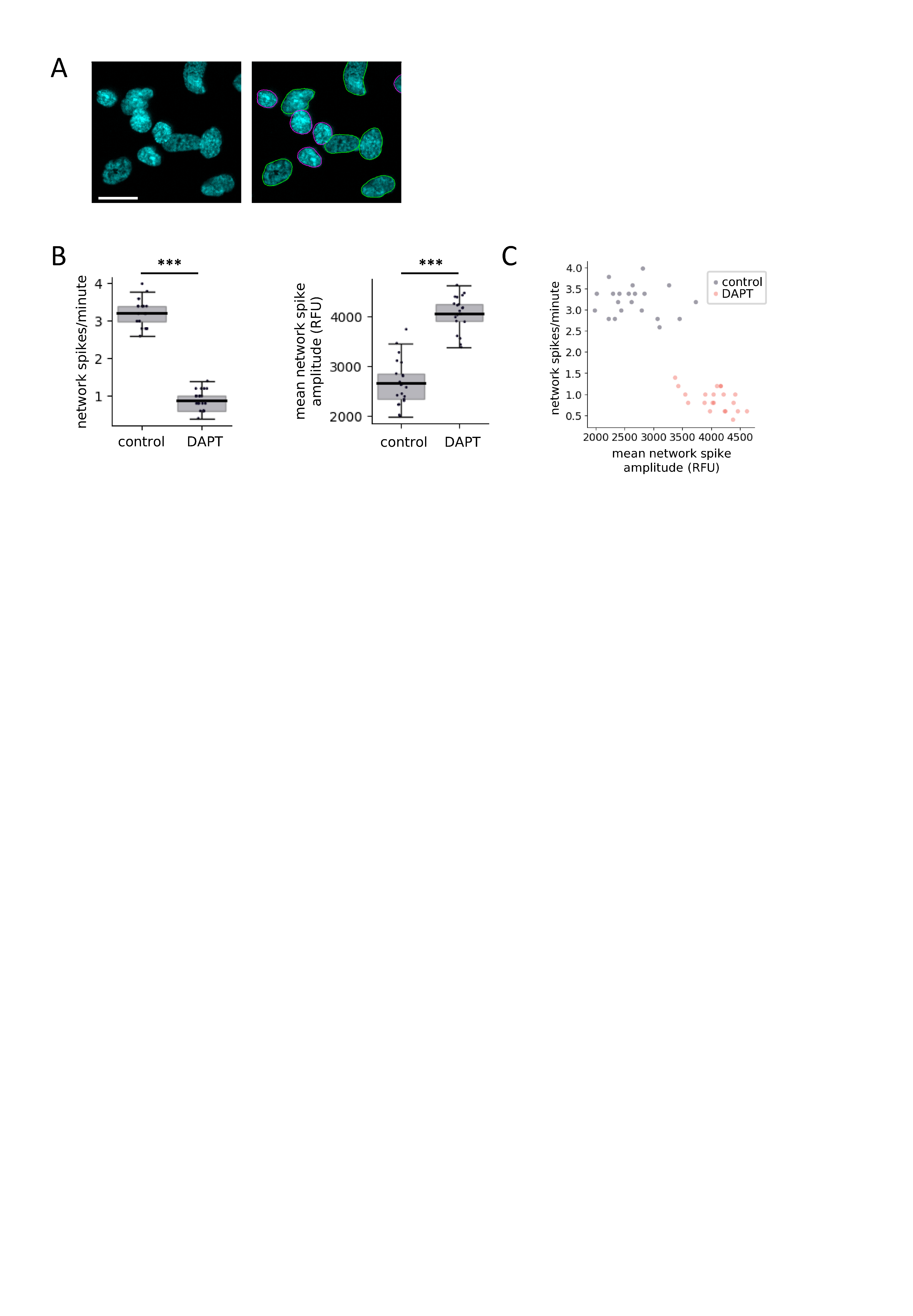

### Figure S4

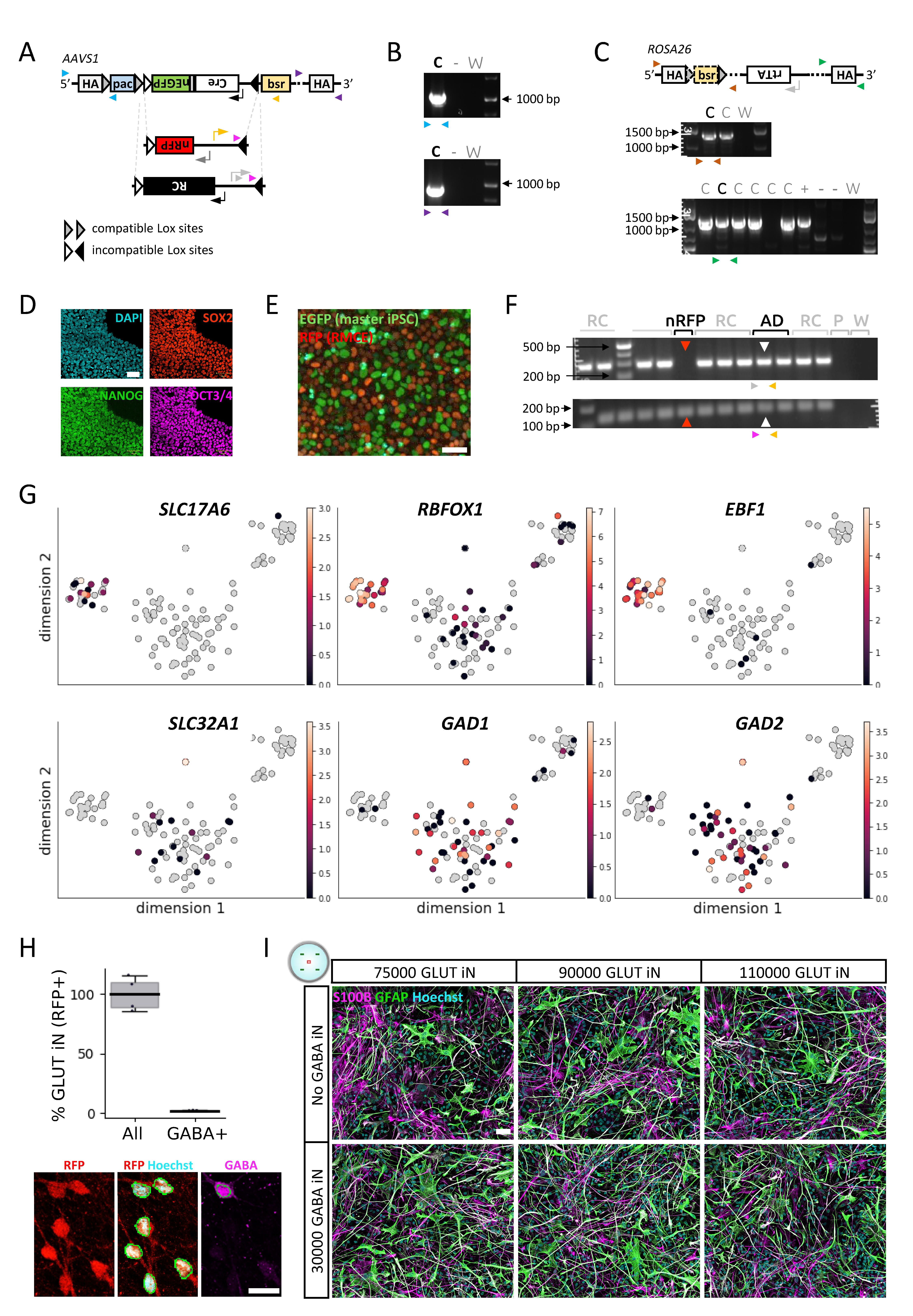

### Figure S5

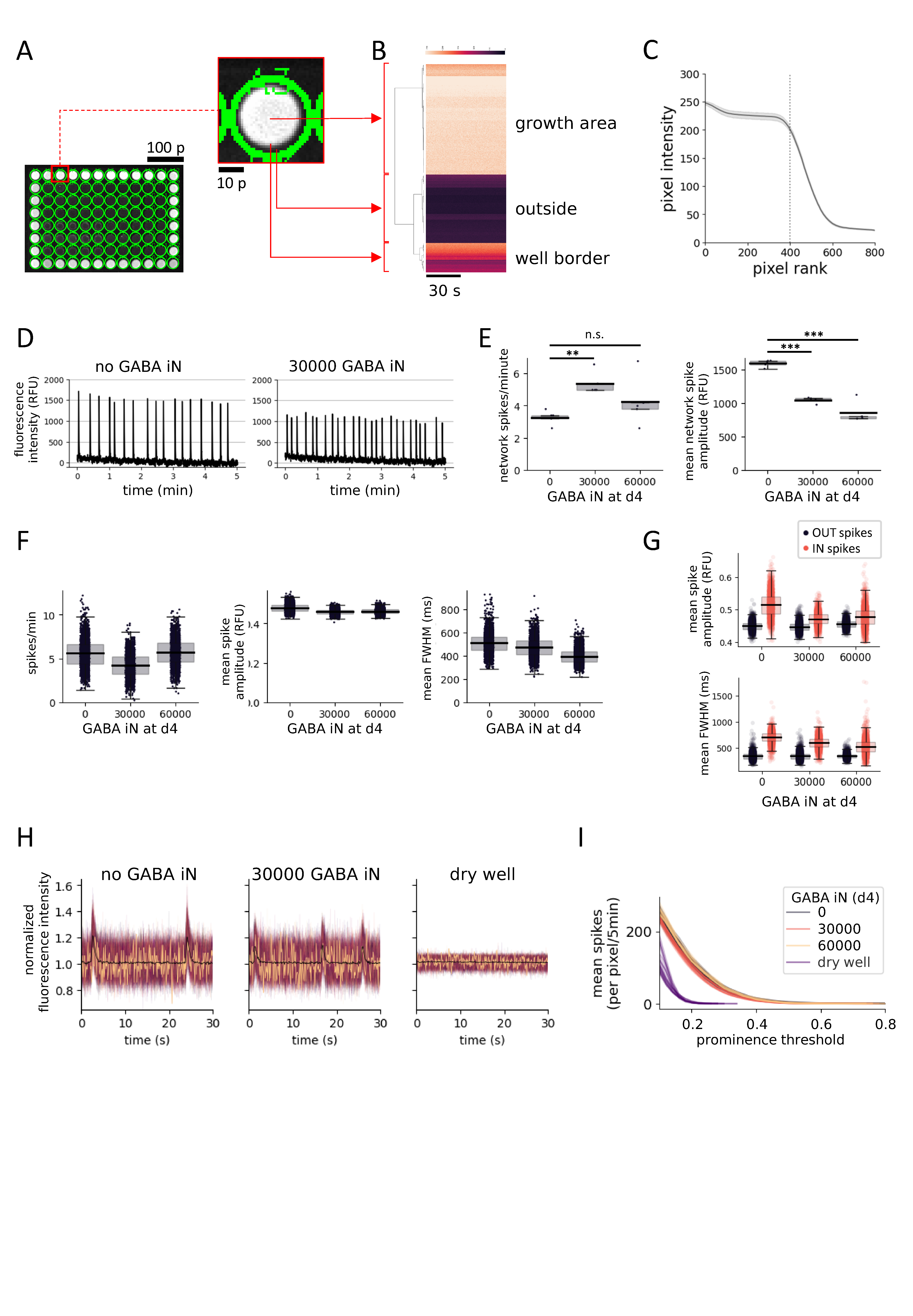

### Figure S6

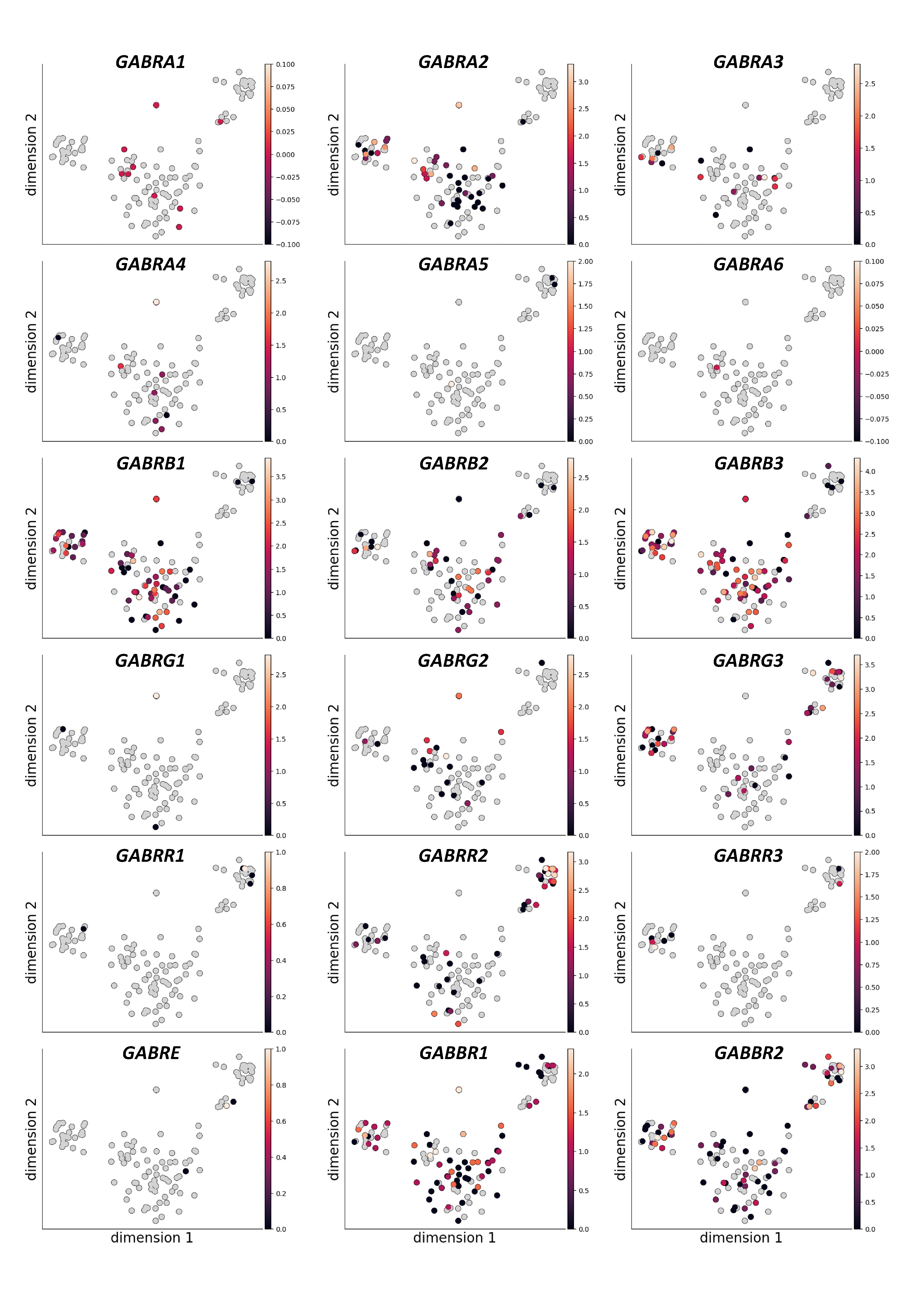

### Figure S7

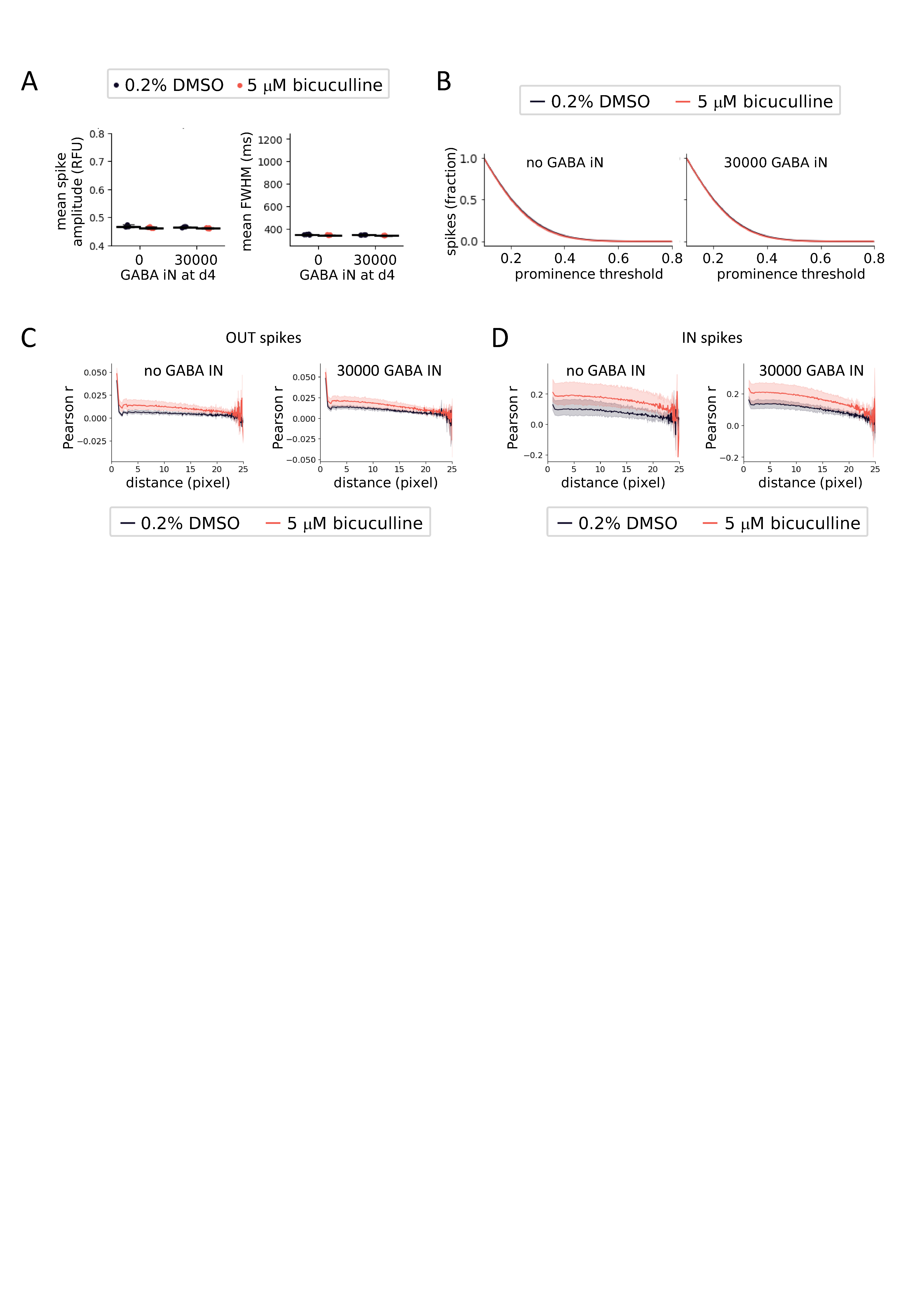
