## Supplementary material for "Development and characterization of a scalable calcium imaging assay using human iPSC-derived neurons": Table S1

| Reagent or Resource | Source | Identifier |
|---|---|---|
| **Antibodies** | | |
| Anti-SOX9 1:500 | Sigma-Aldrich | AB5535 |
| Anti-Gaba 1:100 | Sigma-Aldrich | A2052 |
| Anti-Rabbit IgG (H+L), F(ab')2 Fragment (Alexa Fluor® 488 Conjugate) | Cell Signaling | 4412 |
| Goat anti-Rabbit IgG (H+L) Alexa Fluor™ 647 | Invitrogen | A21244 |
| DLX2 Recombinant Rabbit Monoclonal Antibody (1H13L11) | ThermoFisher | 702009 |
| Anti-GAD67 Antibody, CHEMICON®, mouse monoclonal, 1G10.2 | Merck | MAB5406 |
| Goat anti-Mouse IgG1 Cross-Adsorbed Secondary Antibody, Alexa Fluor™ 647 | Invitrogen | A-21240 |
| **Chemicals and peptides** | | |
| mTeSR Plus | STEMCELL | 100–0276 |
| Geltrex™ Reduced-Growth Factor Basement-Membrane Matrix, LDEV-free, stem-cell qualified | Gibco | A1413302 |
| ReLeSR | STEMCELL | 100-0484 |
| Thiazovivin | Enzo | LKT-T3132 |
| Alt-R™ S.p. HiFi Cas9 Nuclease V3 | IDT | 1081060 |
| CloneR™2 | STEMCELL | 100-0691 |
| Accutase® solution | Sigma-Aldrich | A6964 |
| Alt-R™ HDR Enhancer V1 | IDT | 10181073 |
| Blasticidin S HCl | Gibco | A1113903 |
| Puromycin | InvivoGen | ant-pr-1 |
| Matrigel hESC-qualified, LDEV-free | Corning | 354277 |
| DMEM/F12 (L Glutamine, HEPES, phenol red) | Gibco | 11330032 |
| MEM NEAA (100X) | Gibco | 11140035 |
| hBDNF | Preprotech | 450-02 |
| mouse Laminin | Gibco | 23017-015 |
| hNT-3 | Preprotech | 450-03 |
| Doxycycline | Sigma | D9891 |
| DAPT | Sigma | D5942 |
| Neurobasal-A | Gibco | 10888-022 |
| B-27 supplement 50X | ThermoFisher | 17504001 |
| Bambanker | FujiWakoChemI GZ instruments AG | 302-14681/CS-02-001 |
| Neurobasal-A | Gibco | 10888-022 |
| Neurobasal-Plus | Gibco | A3582901 |
| B27 supplement | Gibco | 17504044 |
| B27 Plus supplement | Gibco | A3582801 |
| GlutaMAX | Gibco | 35050-061 |
| Astrocyte medium | ScienCell | 1801 |
| Ara-C | Sigma-Aldrich | C6645 |
| FBS | ScienCell | 10 |
| Incucyte® Caspase-3/7 Dye | Sartorius | 4440 |
| Gelatin | Sigma-Aldrich | G7041 |
| IGEPAL® CA-630 | Sigma-Aldrich | I3021 |
| KCl | Thermo Scientific Chemicals | 424090010 |
| $MgCl_2$ | Sigma-Aldrich | M8266 |
| UltraPure™ 1 M Tris-HCl Buffer, pH 7.5 | ThermoFisher | 15567027 |
| Tween® 20 | Sigma-Aldrich | P9416 |
| Proteinase K | Roth | 7528.3 |
| BSA Solution 10% | Sigma | A1595 |
| Trizma Hydrochloride Solution | Sigma | T2194 |
| Sodium Chloride Solution, 5M | Sigma | 59222C |
| Magnesium Chloride Solution, 1M | Sigma | M1028 |
| Nonidet P40 Substitute | Sigma | 74385 |
| RNase Inhibitor | ThermoFisher | AM2682 |
| PBS | ThermoFisher | 14190 |
| HEPES | Sigma | H7523 |
| Glucose | Sigma | G7528 |
| DPBS (10X), calcium, magnesium | ThermoFisher | 14080048 |
| Goat Serum | ThermoFisher | 16210064 |
| Hoechst 33342 | ThermoFisher | H3570 |
| Triton X-100 | Sigma-Aldrich | X100 |
| BSA, heat shock fraction, fatty acid free, ≥98% | Sigma-Aldrich | A3803 |
| PFA | Thermo Scientific | 28908 |
| **Compounds** | | |
| (-)-Bicuculline methiodide | Tocris | 2503 |
| 4-Aminopyridine | Sigma-Aldrich | 275875 |
| GABA | Tocris | 0344 |
| Retigabine | Tocris | 6233 |
| **Commercial assays & kits** | | |
| Neon™ Transfection System 10 µL Kit | Invitrogen | MPK1096 |
| Platinum™ Hot Start PCR Master Mix (2X) | Invitrogen | 13001013 |
| Calcium 6 Assay Kit | Molecular Devices | R8191 |
| 3' CellPlex Kit Set A | 10x Genomics | 1000269 |
| GABA ELISA kit | ImmuSmol | BA-E-2500 |
| Fragment Analyzer NGS High Sensitivity Kit | Agilent Technologies | DNF-474-0500 |
| KAPA Library Quantification Kit | Roche Diagnostics | 7960336001 |
| P2 100-cycle kit | Illumina | 20046811 |
| CytoTox 96 non-radioactive assay | Promega | G1780 |
| Incucyte® Caspase-3/7 Green Dye | Sartorius | 4440 |
| Chromium Next GEM Single Cell 3’ Reagent Kit v3.1 | 10x Genomics | PN-1000268 |

| | | |
|---|---|---|
| GABA ELISA kit | ImmusMol | BA-E-2500 |
| **Cell lines and primary cells** | | |
| StemRNA™ Human iPSC 771-3G | Reprocell | RCRP005N |
| BIONi010-C-13 | EBiSC | BIONi010-C-13 |
| Human Astrocytes | ScienCell | 1800 (lot 31978) |
| **CRISPR oligonucleotides** | | |
| crRNA AAVS1-1 protospacer | IDT | ACCCCACAGTGGGGCCACTA |
| crRNA AAVS1-2 protospacer | IDT | GTCACCAATCCTGTCCCTAG |
| crRNA Rosa26-L protospacer | IDT | GTCGAGTCGCTTCTCGATTA |
| crRNA Rosa26-R protospacer | IDT | GGCGATGACGAGATCACGCG |
| Alt-R™ CRISPR-Cas9 tracrRNA, ATTO™ 550 | IDT | 1072533 |
| **Genotyping primers** | | |
| Rosa26_EG2-Fwd | IDT | GTGGGAAGTCGGGAACATAATG |
| Rosa26-EG7R | IDT | CTGGCAACTAGAAGGCACAG |
| pcDNA3.1BGH-rev | IDT | TAGAAGGCACAGTCGAGG |
| Puro F | IDT | GCAACCTCCCCTTCTACGAGC |
| Rosa26-7620R | IDT | ACAGTACAAGCCAGTAATGGAG |
| AAVS1-260F | IDT | TGAGTCCGGACCACTTTGAG |
| AAVS1-1304R | IDT | GTGGGCTTGTACTCGGTCAT |
| AAVS1-6158F | IDT | ACTGTCGGGCGTACACAAAT |
| AAVS1-7108R | IDT | GGC GGA GGA ATA TGT CCC AG |
| AAVS1-1985R | IDT | CTTCCCTGTCCTTGTCGACC |
| Spacer d(ATG)-1F | IDT | GCTGGATTGTAGCTGCTATTAG |
| BSD-4597R | IDT | GAGGGTGGATTCTTCTTGAG |
| CAGGS-5245F | IDT | TCCTACAGCTCCTGGGCAAC |
| **Recombinant DNA** | | |
| pUC_RMCE ROSA26-GCaMP6f-2A-RFP | Azenta | |
| pUC-GW_ROSA26-rtTA | Azenta | |
| pUC-GW_AAVS1-TetO | Azenta | |
| **vessel** | | |
| PhenoPlate 384-well | Revvity | 6057328 |
| PhenoPlate 96-well | Revvity | 6055302 |
