## Supplementary material for "Development and characterization of a scalable calcium imaging assay using human iPSC-derived neurons": Table S2

| compared variables → | | | cell types | | | drug treatment | | | | | |
| --- | --- | --- | --- | --- | --- | --- | --- | --- | --- | --- | --- |
| experiments ↓ | plate format (wells) | coating | ratio GLUT iN/pA | GABA iN | iN/pA nuclei quantification | DAPT | 4-AP | Retigabine | GABA | bicuculline | pixel-resolution analysis |
| 1 | 96 | + | + | - | - | - | + | - | - | - | - |
| 2 | 96 | + | + | - | + | - | + | - | - | - | - |
| 2 | 384 | + | + | - | + | - | + | - | - | - | - |
| 3 | 384 | - | + | + | + | + | - | - | + | - | - |
| 4 | 384 | - | - | + | + | + | + | + | + | - | - |
| 5 | 384 | - | - | - | + | + | + | + | - | - | - |
| 6 | 384 | - | - | - | - | + | + | + | + | - | - |
| 9 | 96 | - | - | + | + | + | - | - | + | - | + |
| 10 | 96 | - | - | + | + | - | - | - | - | + | + |
| 11 | 96 | - | - | - | + | - | + | + | + | - | - |
| 12 | 96 | - | + | + | + | - | - | - | - | + | + |
| 13 | 96 | - | + | + | + | - | + | - | - | + | + |
| Total N | | 3 | 6 | 6 | 10 | 5 | 8 | 4 | 5 | 3 | 4 |
